# Pathogen and pest risks to vegetatively propagated crops in humanitarian contexts: Toward a national plant health risk analysis for Cameroon and Ethiopia

**DOI:** 10.1101/2024.02.12.580019

**Authors:** Romaric A. Mouafo-Tchinda, Berea A. Etherton, Aaron I. Plex Sulá, Jacobo Robledo, Jorge Andrade-Piedra, Kwame Ogero, Bonaventure A. Omondi, Margaret A. McEwan, Paul M. Tene Tayo, Dieudonné Harahagazwe, Mihiretu Cherinet, Setegn Gebeyehu, Louise Sperling, Karen A. Garrett

## Abstract

In humanitarian contexts, supporting agricultural recovery after natural or human-driven shocks—and strengthening vulnerable communities through development interventions—is critical when crop and seed shortages threaten food security and livelihoods. Cameroon and Ethiopia face humanitarian crises that disrupt the production of key food security crops like banana and plantain, cassava, potato, and sweetpotato. In this study, we address crop pathogen and pest effects in a humanitarian context, both in general and with a focus on their effects on planting material quality through seed degeneration. We provide the foundation for a ‘national plant health risk analysis’ to inform regional and national surveillance and management strategies. We analyzed cropland density maps and applied network centrality metrics to identify locations that are candidates for disease and pest management and surveillance in Cameroon and Ethiopia. We used expert knowledge elicitation and global trade data to map pathogen and pest movement risks and characterize trade of crop-specific commodities and planting materials. We identified specific locations that may be important for pathogen and pest spread in Cameroon and Ethiopia, given their roles in the cropland network, and the reported presence of pathogens and pests. These results provide a baseline national plant health risk analysis for Cameroon and Ethiopia. We discuss strategies for ongoing improvement of a national plant health risk analysis, to inform decision-making in the humanitarian sector for designing on-the-ground actions to avoid unintentional spread of pathogens and pests during agricultural recovery interventions.

## 1 Introduction

### 1.1 The importance of root, tuber, and banana crops for low-income countries and challenges for crop health

In the face of increasing natural and human-driven disasters, staple food crops like roots, tubers, and bananas (RTBs) are essential for food security (Nanbol & Namo, 2019; Petsakos et al., 2019; Scott, 2021; Garrett et al., 2022). RTBs are vegetatively propagated, with planting material (‘seed’) that includes suckers (bananas and plantains, (*Musa* spp.)), stakes (cassava), tubers (potato), and vines (sweetpotato). Foundational studies such as Fauquet and Fargette (1990), Thresh and Otim-Nape (1994), and Otim-Nape and Thresh (1998) developed understanding of the spread and management of cassava mosaic viruses in Africa. Vegetative propagation increases the risk of seed degeneration (the build-up of plant pathogens and pests and pathogens in planting material), facilitating the unintentional spread of pests and pathogens through seed trade or replanting, particularly in informal trade systems (Andrade-Piedra et al., 2025; Gibson & Kreuze, 2015; Jeger et al., 2007; Tessema & Tesfaye, 2023; Thomas-Sharma et al., 2016). Informally traded seed of RTB crops (i.e., planting material obtained from neighbors, local markets, or uncertified sources) along with on-farm saved seed, is the primary seed source for growers in low-income countries and often has little or no official phytosanitary control (Gibson & Kreuze, 2015; Jeger et al., 2007; Sperling et al., 2020; Thomas-Sharma et al., 2016). This contributes to ongoing crop production challenges due to RTB diseases and pests. Note that, although this article focuses on humanitarian contexts, throughout the text we use ‘disease’ to refer to crop disease.

Cameroon and Ethiopia rely on informal seed trade and on RTB crops for income generation and food security (Hirpa et al., 2010; OCHA, 2023a). By mapping national and cross-border movement of seed, we identify regions that may be at risk for pathogen spread, through network-based epidemiological modeling. Trade flows can be disrupted or redirected during and after disasters through population displacement, market collapse, and emergency seed distribution (Gilligan, 2008; Jeger et al., 2004; Jeger et al., 2007). The analyses here are part of an ongoing ‘national plant health risk analysis’ proposed for these crops and implemented as proof of concept in this paper.

### 1.2 Ongoing disasters and humanitarian needs in Cameroon and Ethiopia

Disaster plant pathology addresses how disasters influence plant disease dynamics, through human migration, infrastructure destruction, aid distribution, and creation of more suitable environments for plant diseases (Etherton et al., 2024). Disasters like droughts, floods, forest fires and hurricanes can directly impact plant health, through the spread of pathogens and pests, especially impactful if the species are new introductions to the region. Political unrest, conflicts, and wars can indirectly affect crop health, through damaged infrastructure and reduced capacity. Disasters can trigger epidemics and exacerbate food insecurity in vulnerable communities (Etherton et al., 2024).

Humanitarian challenges in Cameroon and Ethiopia motivate the analysis of these countries’ crop health risks and seed systems in the context of disaster plant pathology (Etherton et al., 2024). In Cameroon, crises including natural disasters and armed conflicts have displaced growers and disrupted agricultural production (Bang, 2021; Bang & Balgah, 2022; Bang & Balgah, 2022; Mbah et al., 2023; FAO, 2021a, 2021b; OCHA, 2023a) (**Figure 1**). For example, Cameroon hosts more than 450,000 refugees and asylum seekers annually, and the demand for humanitarian aid continues to grow (Nisbet et al., 2022.Macrotrends, 2023; OCHA, 2022; Foyou et al., 2018). However, humanitarian access remains a major challenge due to administrative obstacles, insecurity due to armed groups, and infrastructure damage (ECPHAO, 2023).

**Figure 1.**
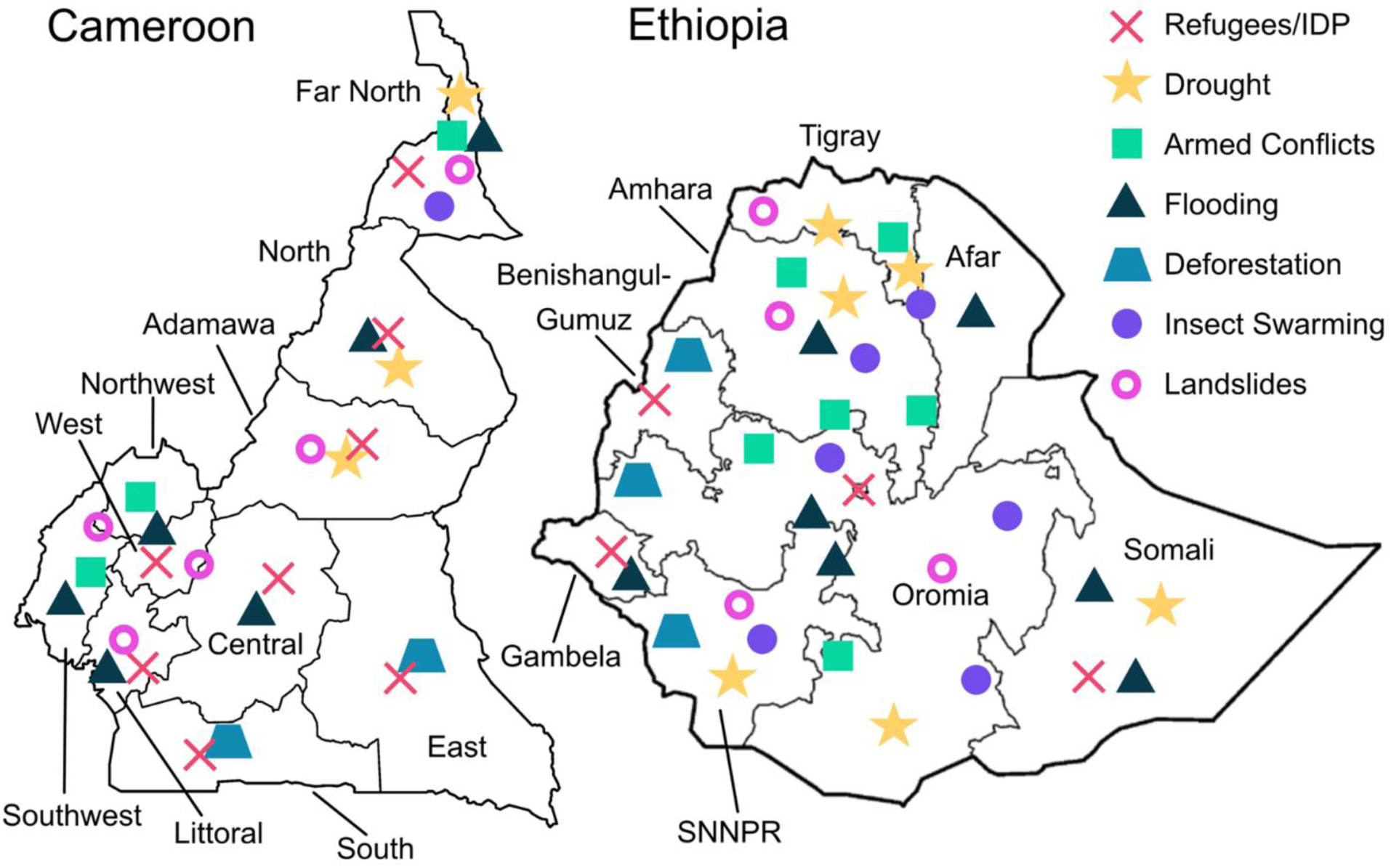
Humanitarian crises and disaster hazards affecting agricultural production in Cameroon and Ethiopia. Symbols indicate representative categories of disruption (see legend). Locations are approximate and illustrative, not comprehensive. Adapted from Bang and Balgah (2022), IOM (2023), and Raleigh et al. (2010). Ethiopia boundaries follow 2022; SNNPR = former Southern Nations, Nationalities, and Peoples’ Region.

Ethiopia faces drought, floods, armed conflicts, pest outbreaks, and political instability, of which drought is a main factor reducing agricultural production and increasing malnutrition among the affected communities (Alasow et al., 2023; CDP, 2023; Wubaye et al., 2023) (**Figure 1**). Ethiopia has experienced armed conflicts such as the Tigray civil conflict. The compound effects of climate shocks, armed conflicts, and displacement, continue to drive humanitarian needs, and more than 20 million people required food assistance in 2023 (OCHA, 2023b). Many disaster-affected growers cannot save seed, creating demand for emergency seed supply alongside inputs and cash; Ethiopian farmers have received such aid for over 40 years (Bramel et al., 2004; Man, 2018; Mengistu, 2001).

### 1.3 The need for analysis of crop health risk to inform humanitarian aid interventions

The humanitarian imperative goes beyond saving lives and addressing immediate needs; it also aims to foster livelihood recovery and resilience (Hunt et al., 2018; USAID, 2022). We use ‘humanitarian contexts’ to include both post-crisis agricultural recovery and also development interventions in vulnerable communities, in contrast to settings where food production and availability are stable at acceptable levels (Sperling, 2008). A key priority is ensuring access to pathogen- and pest-free seed, including quality seed of crop varieties and seed types that meet farmer preferences (Kwambai et al., 2024; Nduwimana et al., 2022; Singh et al., 2015; Sperling et al., 2022). Seed-borne threats can be amplified during crises because seed aid may arrive too late, provide unsuitable varieties, or inadvertently introduce new pests or diseases (Sperling & McGuire, 2009).

For organizations planning assistance, there is often limited information available to assemble a ‘national plant health risk analysis’ that includes the range of pathogens and pests that threaten crop health, along with their potential spread through planting materials and other means. In both Cameroon and Ethiopia, research has documented major diseases affecting RTB crops, such as banana bunchy top disease, cassava mosaic diseases, and bacterial wilt (Doungous et al., 2022; Ngatat et al., 2024; Poubom et al., 2005; Tessema & Seid, 2023) as well as key pests including aphids, whiteflies (*Bemisia tabaci*), and weevil (including *Cosmopolites sordidus*, *Cylas puncticollis,* and *C. brunneus*). However, these studies primarily focus on individual pathogens and pests, and there is a lack of research assessing the broader risk of geographic spread in these regions (Szyniszewska et al., 2021). The common lack of systematic and current pathogen and pest information, that could support a national plant health risk analysis, highlights the importance of rapid risk assessment tools that identify regions likely to be important in pathogen and pest spread, target economically important pathogens and pests, and assess the potential role of informal trade in pathogen and pest spread (Carvajal-Yepes et al., 2019; Thomas-Sharma et al., 2017; Xing et al., 2020). This information can help in identifying risks for consideration in emergency seed assistance and planning longer-term agricultural development strategies.

### 1.4 The objectives and approaches of this study

This paper lays the foundation for a national crop health risk analysis to support decision-making about crop protection and healthy seed systems. Although a national plant health risk analysis is broadly valuable for plant health management under any circumstances, it takes on particular urgency in humanitarian contexts where seed systems are disrupted, phytosanitary oversight is weakened, and emergency seed distribution risks unintentional pathogen and pest spread. We use two complementary approaches. The first is cropland connectivity analysis, which provides maps of the likely role of geographic locations in the spread of pathogens and pests (Plex Sulá et al., 2025; Margosian et al., 2009; Xing et al., 2020; Andersen Onofre et al. 2021; Buddenhagen et al. 2022). The second is expert knowledge elicitation, which can provide information about crop health based on experts’ knowledge and identifies gaps in domain knowledge (Andersen Onofre et al., 2021; Thomas-Sharma et al., 2017; Hughes and Madden, 2002; Robledo et al., 2025). Expert knowledge has biases and other limitations, but expert knowledge elicitation is an efficient approach to systematically generate a summary of current understanding, as demonstrated across a range of disciplines (Arndt et al., 2022; Battiston et al., 2021; Chen et al., 2019; EFSA, 2014; EFSA Panel on Plant Health, 2018; Hadjigeorgiou et al., 2022; Knol et al., 2010; Motisi et al., 2022; Soares et al., 2024; U.S. Environmental Protection Agency, 2009). Together these approaches provide rapid risk assessments and a baseline for follow-up studies as resources allow.

The objectives of this study are to 1) identify locations in Cameroon and Ethiopia likely to be important for pathogen and pest spread in potato and sweetpotato, given the locations’ roles in each country’s network of crop host availability, 2) evaluate disease and pest risk in Cameroon and Ethiopia using expert knowledge elicitation, and 3) identify potential risks to the spread of pathogens and pests through informal seed trade and food commodity trade. We expand these analyses for Cameroon to include bananas (by which we generally mean bananas and plantains) and cassava. These objectives are pursued with a particular focus on informing decision-making in humanitarian contexts, where the results can guide seed sourcing, surveillance priorities, and phytosanitary measures during agricultural recovery interventions. We provide a framework in early TRL of a ‘national plant health risk analysis’ for these RTB crops in Cameroon and Ethiopia that generates actionable insights for humanitarian organizations, growers, RTB researchers, and national programs, with the potential to inform regulatory policies and guide on-the-ground practical actions.

## 2 Methods

The analyses were implemented with code available in GitHub at https://githu.com/GarrettLab/Mouafo-Tchinda-Etherton-et-al-2024. Versions of these analyses are part of the ongoing development of the R2M Plant Health Toolbox for Rapid Risk assessment to support Mitigation of plant pathogens and pests (garrettlab.com/r2m/). The current study was built on insights gained from our previous capacity needs assessment surveys among humanitarian organizations in Cameroon, the Democratic Republic of Congo, Ethiopia, Haiti, Mozambique, Bangladesh, and Madagascar (Andrade-Piedra et al., 2023; Canney-Davison et al., 2023).

### 2.1 Cropland connectivity for host-specific pathogens and pests

We used a cropland connectivity analysis to map locations likely to be important in outbreaks of pathogens and pests of bananas and plantains (henceforth, ‘banana’), cassava, potato and sweetpotato. We used crop-specific harvested area maps (raster files) for banana, cassava, potato and sweetpotato (International Food Policy Research, 2020), where each pixel represents the amount of cropland (in ha) that was harvested in 2017. Cameroon and Ethiopia were the focus countries in these analyses, where the cropland within each country and in the immediate surrounding countries (±0.5° latitude and longitude) were included. Locations (or individual cells within the cropland raster maps) with harvested areas greater than zero were retained as network “nodes,” and a gravity model was used to evaluate the relative likelihood of dispersal between each pair of nodes (Jongejans et al., 2015). We used the 2017 harvested-area rasters as a proxy for host availability to support rapid geographic prioritization, recognizing that humanitarian disruptions can shift cropping patterns over time. In the gravity model, the ‘mass’ of each node is the harvested area of the crop in that pixel (raster cell), so nodes with larger harvested area have greater pull in the dispersal network. Thus, in epidemiological applications, the gravity model reflects how greater host availability at a node can make it a more important inoculum source and also make inoculum that reaches the node more likely to reach the host.

In separate analyses for each crop, the importance of each node in epidemic/invasion networks was evaluated using a cropland connectivity risk index (CCRI) as described by Xing et al. (2020), using the geohabnet package in R (version 4.3.2; Plex Sulá et al., 2025). The CCRI was defined as the weighted sum of four network metrics, to capture a range of potential roles a location may have during an epidemic or pest invasion. These network metrics include 1) betweenness centrality, indicating a node’s role as a pathway between other nodes in a network (bridge nodes), 2) node strength, indicating its connection to other nodes in the cropland network, 3) the sum of a node’s nearest neighbors’ node degrees, indicating how connected a node’s closest neighbors are, and 4) eigenvector centrality, which accounts for both quantity and quality of links in the cropland network. Higher CCRI scores indicate locations that are more likely to play an important role in an epidemic or invasion based on host availability and pathogen or pest dispersal, according to their geographic proximity and likely role in an epidemic or invasion network.

### 2.2 Expert knowledge elicitation

We developed surveys for use in expert knowledge elicitation for Cameroon and Ethiopia to capture the knowledge of RTB experts currently working in the field. Considering the growing but still limited databases available to support seed system responses in humanitarian situations (see https://seedsystem.org/field-assessments-action-plans/), and to support general strategies for RTB crop health, the goal was to assemble current expert knowledge in a useful framework to support decision-making. Although expert knowledge elicitation has limitations, including potential knowledge gaps due to expert experience, personal biases, and data availability, it remains a valuable tool for risk assessment (Robledo et al., 2025). It can be used to assess the risk of seed-borne pest and pathogen spread in vulnerable regions, providing critical insights where empirical data may be lacking. To reduce these limitations, elicitation protocols offer strategies to address how expert judgments are structured, shared, and aggregated (e.g., behavioral approaches vs. formal weighting/aggregation frameworks) (Cooke, 1991; Colson & Cooke, 2018; Hemming et al., 2018). Expert knowledge summaries can help stakeholders anticipate and mitigate threats, and guide the development of management strategies against emerging plant pathogens and pests before extensive field studies.

The first stage of survey development was to review the content of the country- and crop-specific surveys with the organizing team to make sure they were appropriately matched to the situation in each country. The surveys focused on RTB pathogens and pests causing high yield loss, with potentially important impacts on vulnerable human populations. In Cameroon, four surveys covered bananas, cassava, potato and sweetpotato. Similarly, two surveys were designed for experts in Ethiopia, focusing on potato and sweetpotato. Surveys were deployed using Google Forms (Vasantha Raju & Harinarayana, 2016). Crop-specific experts who had insights into seed systems participated in one-day workshops in November 2022 in Yaoundé, Cameroon, and in March 2023 in Addis Ababa, Ethiopia. In Cameroon, we recruited 18 experts for cassava, 13 for potato, 10 for banana and 7 for sweetpotato. In Ethiopia, we recruited 20 experts for potato, and 9 for sweetpotato. Experts known through previous collaboration with CGIAR and the University of Florida were selected based on their experience with RTB crops, representing governmental organizations, the private sector, NGOs, National Agricultural Research and Extension Systems (NARES), and CGIAR centers (IITA, CIP, and Alliance Biodiversity-CIAT). Across crops and locations, 60 to 78% of participating experts identified as seed system experts in Cameroon or seed system/breeding experts in Ethiopia. Anonymized information about the experts’ experience in each state or region and each crop, along with all crop-specific responses, are available upon request. Participating experts completed the surveys individually, for the crop for which they had expertise. Experts were first asked to provide their years of experience in each region or state in their country of expertise. In each survey, experts were then asked to identify the major pests and diseases, and identify the informal exchange routes for seed in the country and with neighboring countries, estimating crop-system specific parameters. Because not all major pests and pathogens are seed-borne, experts also separately identified those they considered key contributors to seed degeneration, which are highlighted in the crop-specific results. Targeted questions in the expert knowledge elicitation in Ethiopia were also used to parameterize models used in a study of the geographic distribution of pathogen risk for potato in Ethiopia (Etherton et al., 2025).

Answers from experts were compiled and organized by administrative regions or states in each country. In Ethiopia, state boundaries have now changed, but the regional states in March 2023 were Tigray, Afar, Amhara, Benishangul-Gumuz, Somali, Oromia, Gambela, Southwestern, Southern Nations, Addis Ababa, Dire Dawa and Harari. An initial list of key pathogens and pests was drawn from previous studies carried out in both countries, and based on preliminary discussions with the organizing team. The pathogens and pests reported in the expert knowledge elicitation to be present in regions or states were compiled, and visualized using polar bar plots using the R package ggplot2 (version 3.3.6; Wickham, 2016). Experts were not asked to identify the absence of pests and pathogens, so a lack of reports on a pest or pathogen’s presence in a region or state does not necessarily imply absence, only that experts were not aware of its presence. We used the log of the sum of the total years of experience reported by experts for each region or state as a proxy for relative confidence in the reports for that region or state. For both countries, pest and pathogen community maps were created for potato and sweetpotato, and for Cameroon also for cassava and banana. Note that an important caveat for comparison of locations is that where there is less crop production there tends to be less attention from experts, so lower expert experience in a region in a sense represents ‘lower sampling effort’. That is, regions and regional states where experts have less experience will tend to have lower numbers of pathogen and pest species reported. Visualizations for each crop indicate whether at least one expert reported each pest or pathogen in each region or state. To address the sampling effort issue, visualizations also indicate the corresponding relative confidence for the region or state based on the total years of expert experience for the region or state.

### 2.3 Within-country informal seed trade in Cameroon

Experts in Cameroon were asked about informal seed trade of potato, sweetpotato, cassava, and bananas between regions. In the expert knowledge elicitation, informal seed trade was defined as ‘seed obtained from neighbors, local markets, or uncertified seed producers or traders. For each potential link between a pair of regions, the link weight assigned was the proportion of experts who reported that there was informal seed trade between that pair. This proportion was used as a measure of confidence for the potential existence of seed trade between two regions or states. Higher awareness of trade among experts may be associated with higher trade volume. The node strength, or the sum of link weights, of informal seed trade was then calculated for each region using the igraph R package (version 1.5.0; Csárdi et al., 2023). These estimates of informal trade were compiled for each crop in Cameroon and represented as networks using the ggraph R package (version 2.1.0; Pedersen, 2022). The informal seed trade networks between regions were then plotted on a map of Cameroon with political administrative boundaries, using the rgeoboundaries R package (version 1.2.9000; Dicko, 2023; Runfola et al., 2020).

### 2.4 Trade with neighboring countries: Informal seed trade and food commodity trade

Experts were asked about informal trade of potato, sweetpotato, cassava and banana seed with neighboring countries. Experts indicated which countries they knew informally imported seed from and exported seeds to Cameroon or Ethiopia for each crop, where “informal seed” was defined as seed “obtained from personal seed (seed saved from prior harvests/stocks), neighbors, local markets, or uncertified seed producers.” We created an informal trade network based on the total number of experts who reported that trade occurred between two countries. We rescaled these reports to represent the proportion of experts who identified potential informal trade routes, for example, if 7 of a total of 10 experts identified trade between Cameroon and the Central African Republic, the link weight would be 0.7. We created these informal import and export networks for potato, sweetpotato, cassava, and banana for Cameroon, and potato and sweetpotato for Ethiopia, and represented them as networks using the ggraph R package (version 2.1.0; Pedersen, 2022).

Trade in food commodities generally represents a lower risk for the spread of pathogens and pests compared to trade in planting materials, but can also be a factor if effective phytosanitary measures are not in place. For crops like potato trade in food commodities is a higher risk, because tubers that are imported as food may later be used as seed; planting of food commodities is more unusual for sweetpotato, and not a risk for cassava and banana. For the analysis of food commodity trade, FAO import and export data were analyzed for Cameroon, Ethiopia, and their respective neighboring countries (FAOSTAT, 2021). Here, the FAO data represents the trade of food, or “trade that occurs within a legal framework and is obtained from certified governmental distributors.” Products manufactured by roasting or sterilization were excluded in this trade data, because of their reduced risk. FAO trade data were assembled for each crop, where the trade volume (in tons) between each pair of countries was averaged for 2006-2020 (FAOSTAT, 2021). The trade volumes were then rescaled to represent the proportion of crop-specific trade for a country. For example, if Ethiopia exported on average 300 kg of potato to Djibouti, where the most exported was 1000 kg, the link weight would be 0.3. The FAO food trade data were assembled in a network and compared with the informal seed trade as reported by experts.

## 3 Results

### 3.1 Cropland connectivity

In Cameroon, the analysis of crop landscape connectivity indicated that, for banana pathogens and pests, locations in the Southwest, West, Northwest, Centre, South, East, and Adamawa regions have high banana cropland connectivity (**Figure 2**). For cassava, many regions of Cameroon, except in the Southwest region, have locations with high cassava cropland connectivity within country and to highly connected large cassava croplands in neighboring countries like Nigeria and Chad (**Figure 2**). For potato, locations in the West, Northwest, and Adamawa regions have higher cropland connectivity (**Figure 2**). These higher-risk potato cropping areas are fairly small, reflecting the limited extent of potato production across Cameroon and the geographic isolation from more extensive potato production in neighboring countries. For sweetpotato, locations in the West, Northwest, Adamawa, North and Far-Noth regions have high connectivity (**Figure 2**), based on host availability, with locations in the Northwest and West having the highest cropland connectivity across these regions.

**Figure 2.**
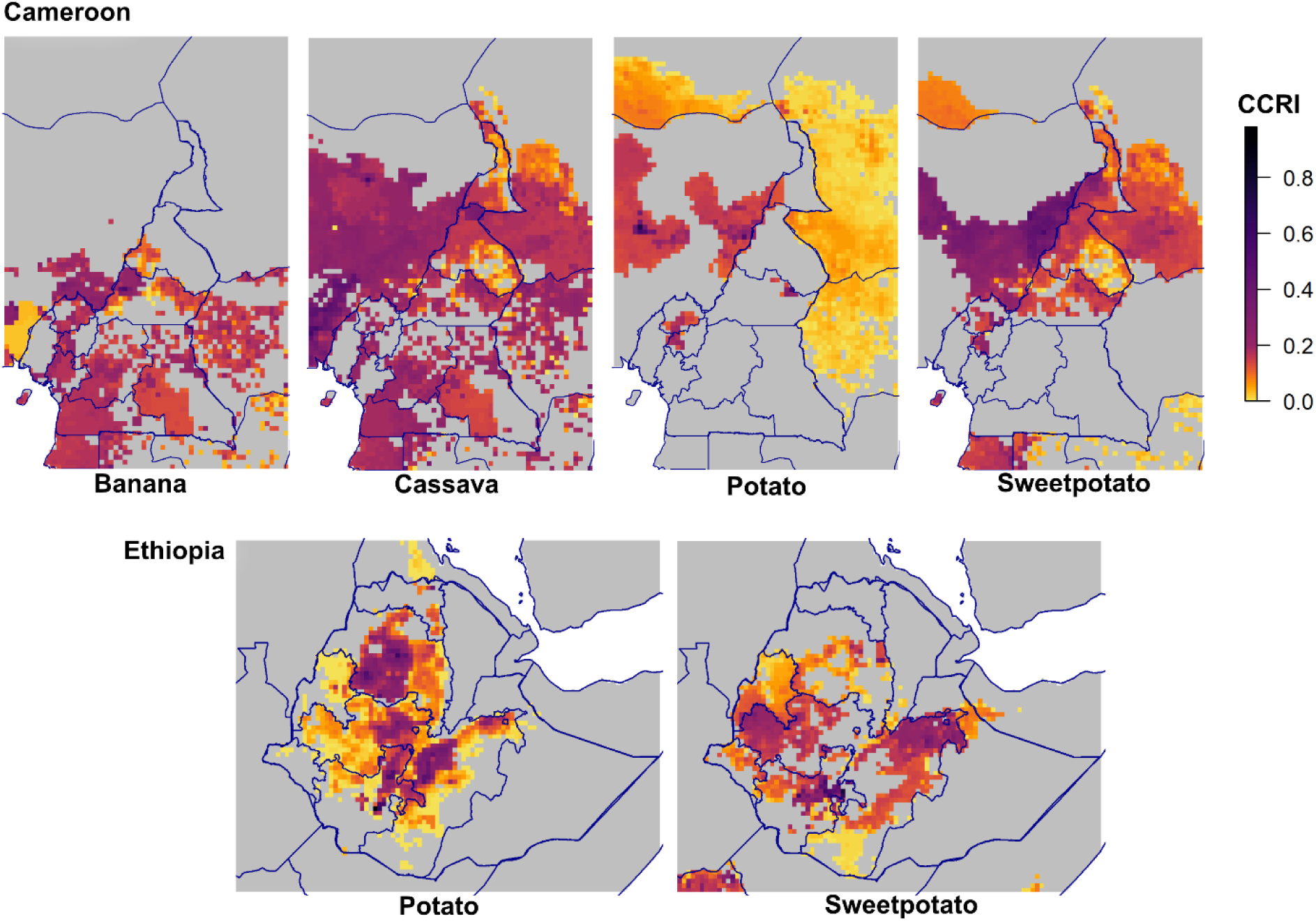
Cropland connectivity identifies likely hub and bridge areas for pathogen and pest spread in RTB crops. Maps show the cropland connectivity risk index (CCRI) for banana, cassava, potato, and sweetpotato in Cameroon and for potato and sweetpotato in Ethiopia. CCRI was calculated from crop harvested-area rasters by treating cropped pixels as network nodes and estimating link strength between nodes with a distance-based gravity model; higher CCRI values indicate locations expected to contribute more to establishment and spread given host availability and landscape connectivity.

For Ethiopian potato production, locations in a large expanse across central Ethiopia have higher cropland connectivity, spanning across Amhara, Oromia, Sidama and into the former Southern Nations, Nationalities and Peoples Region (SNNPR), with the highest CCRI scores occurring in the SNNPR and central Amhara regions (**Figure 2**). For Ethiopian sweetpotato, key locations include parts of western Oromia and Benishangul-Gumuz, the border between the former SNNPR and Sidama, and eastern Oromia including Harari and Dire Dawa (**Figure 2**). The highest CCRI scores for Ethiopian sweetpotato were on the border between Sidama and the former SNNPR. Neighboring countries were not strongly connected to Ethiopia in potato and sweetpotato cropland networks, as the reported harvested cropland production in neighboring countries was relatively low (**Figure 2**) (IFPRI, 2020).

### 3.2 Expert knowledge elicitation: Pathogen and pest communities

Experts noted locations where they knew pathogens and pests were present. Here, absence of evidence is not evidence of absence, but locations noted by experts are generally more well-known to have important problems with pathogens and pests. For bananas in Cameroon, experts had the most experience in the Centre region, followed by the South, Littoral, and then the East and Northwest regions.

The most pathogens and pests, eight, were noted by experts in the South zone. Anthracnose (caused by *Colletotrichum musae*) was noted present in most regions of Cameroon, and nematodes (*Radopholus similis, Pratylenchus goodeyi*) and yellow sigatoka (caused by *Pseudocercospora musae*) were noted present in many regions. Banana seed degeneration in Cameroon was noted by experts to be associated with many pathogens/diseases and pests, including nematodes, BBTV, banana weevil (*Cosmopolites sordidus*), anthracnose, and rhizome soft rot (caused by *Pectobacterium*). In follow-up discussions with a subset of experts, black sigatoka disease (caused by *Pseudocercospora fijiensis*) was also highlighted as important.

For cassava in Cameroon, the East, Centre, and South regions were noted to have the presence of all 13 pathogens that were included in the expert knowledge elicitation survey. Cassava green mite (*Mononychellus tanajoa*), cassava root rot (caused by *Fusarium* spp.), cassava mosaic disease, cassava mealybugs (*Phenacoccus manihoti*) and cassava whitefly (*Bemisia tabaci*) were noted as present in every region across Cameroon. Cassava seed degeneration was noted to be associated with many pests and diseases, including CMV, cassava root rot, cassava anthracnose disease (caused by *Colletotrichum gloeosporioides*), ACMV, *Colletotrichum* spp., and cassava whitefly.

For potato in Cameroon, the western border was noted by experts to have most major pathogens and pests evaluated, with the Northwest and Southwest regions noted as having at least eight of those evaluated (**Figure 3**). Early blight (caused by *Alternaria solani*), PVY, potato aphids (*Myzus persicae*) and Fusarium dry rot (caused by *Fusarium* spp.) were noted to be present in seven of the ten regions across Cameroon. Potato seed degeneration was noted to be associated with late blight (caused by *Phytophthora infestans*), PVY, PLRV, bacterial wilt (caused by *Ralstonia solanacearum*), Fusarium dry rot, and nematodes (*Meloidogyne spp*.).

**Figure 3.**
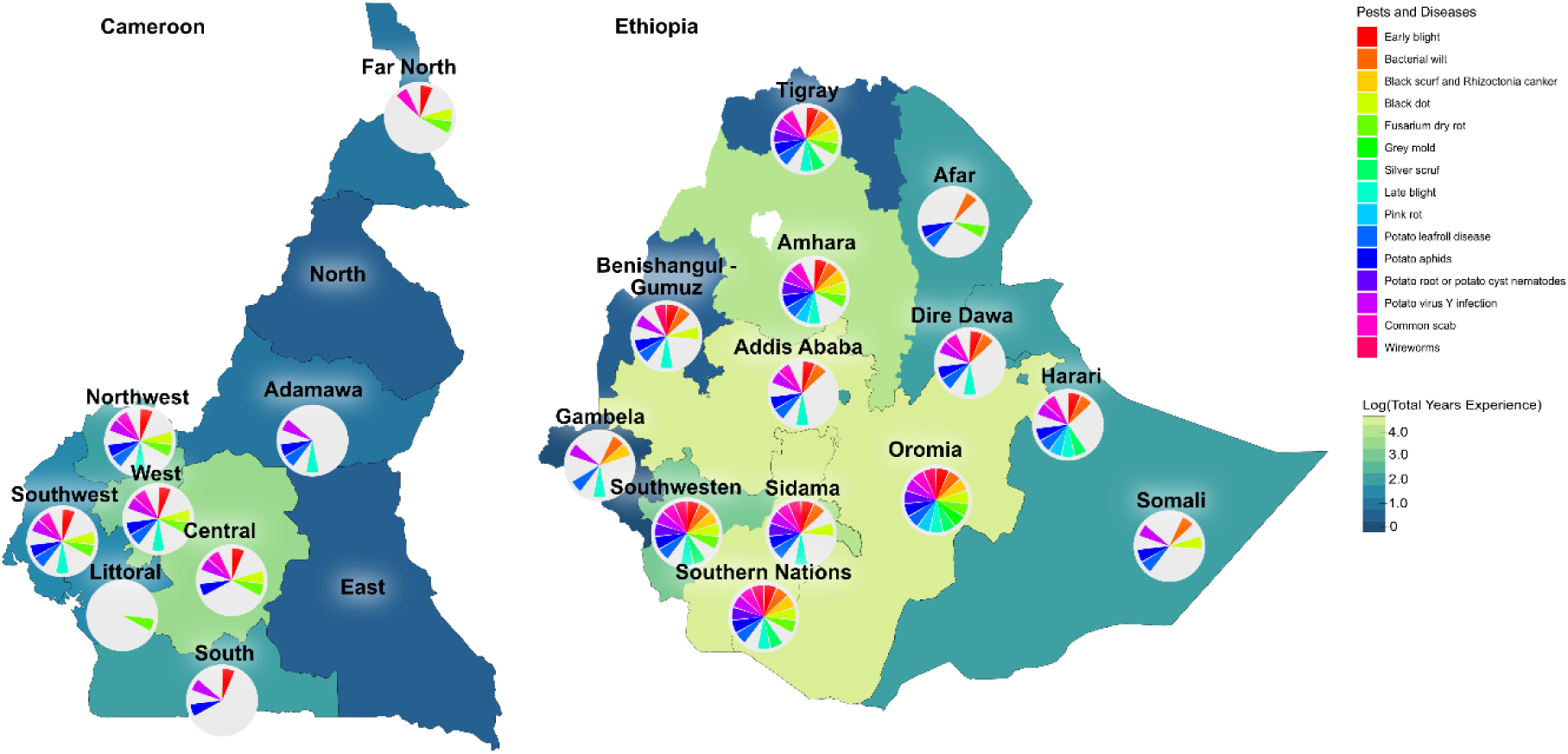
The major potato pests and pathogens noted in expert knowledge elicitation for Cameroon and Ethiopia (based on Ethiopian states as of March 2023). The color in a region or state is proportional to the log-transformed sum of the years of experience reported by the experts for potato in that region or state, and can be interpreted as a measure of confidence in reports by experts for a region or state. The circular bar charts show the pests and diseases reported as present by at least one expert in each region or state.

For sweetpotato in Cameroon, the West region was noted to have most major pathogens and pests, with the South region having six pathogens noted. For the Northwest, West, Centre, and Littoral regions, at least five were noted. Alternaria blight (caused by *Alternaria* spp.) and bacterial wilt (caused by *R. solanacearum*) were noted as present in most regions of Cameroon. Sweetpotato seed degeneration was reported to be associated with sweetpotato viruses, Alternaria blight, and Fusarium root rot. In follow-up discussions with a subset of experts, they noted the importance of weevils (*C. puncticollis* and *C. brunneus*).

For Ethiopian potato, central zones were noted in expert knowledge elicitation to have most major pests and pathogens, with Oromia noted as having at least twelve of the pathogens and pests (**Figure 3**). The former SNNPR, Southwestern, Amhara and Tigray regions each had reported pathogen communities of at least ten pathogens. Experts noted that PLRV was present in every region across Ethiopia. PVY and potato aphids (*M. persicae*) were noted as present in most regions. Experts selected the key causes of potato seed degeneration in Ethiopia to be *Phytophthora infestans*, PVY, PLRV, potato aphids, early blight, and bacterial wilt (caused by *R. solanacearum*).

For Ethiopian sweetpotato, southern zones were noted by experts to have the highest pest and pathogen communities, with Sidama and the former SNNPR both noted as having at least 8 pathogens. Alternaria blight (caused by *A. bataticola*) and the sweetpotato weevil (*C. puncticollis* and *C. brunneus*) were noted as present in most regions across Ethiopia. Experts selected the key causes of seed degeneration as sweetpotato viruses (SPFMV, and SPCSV), sweetpotato weevil, *A. bataticola*, and sweetpotato aphids (*Aphis gossypii* and *M. persicae*).

### 3.3 Expert knowledge elicitation: Within-country informal seed trade in Cameroon

We analyzed the network of seed transfers between regions in Cameroon as reported by experts for potato and sweetpotato, to assess the risk of pathogen and pest spread through seed between regions. For informal potato and sweetpotato seed trade in Cameroon, all the regions were reported to be interconnected. There was high node strength for seed trade reported in the West region for potato (77% of experts noting trade) and in the Centre and Littoral regions for sweetpotato (43% of experts noting trade) (**Figure 4**).

**Figure 4.**
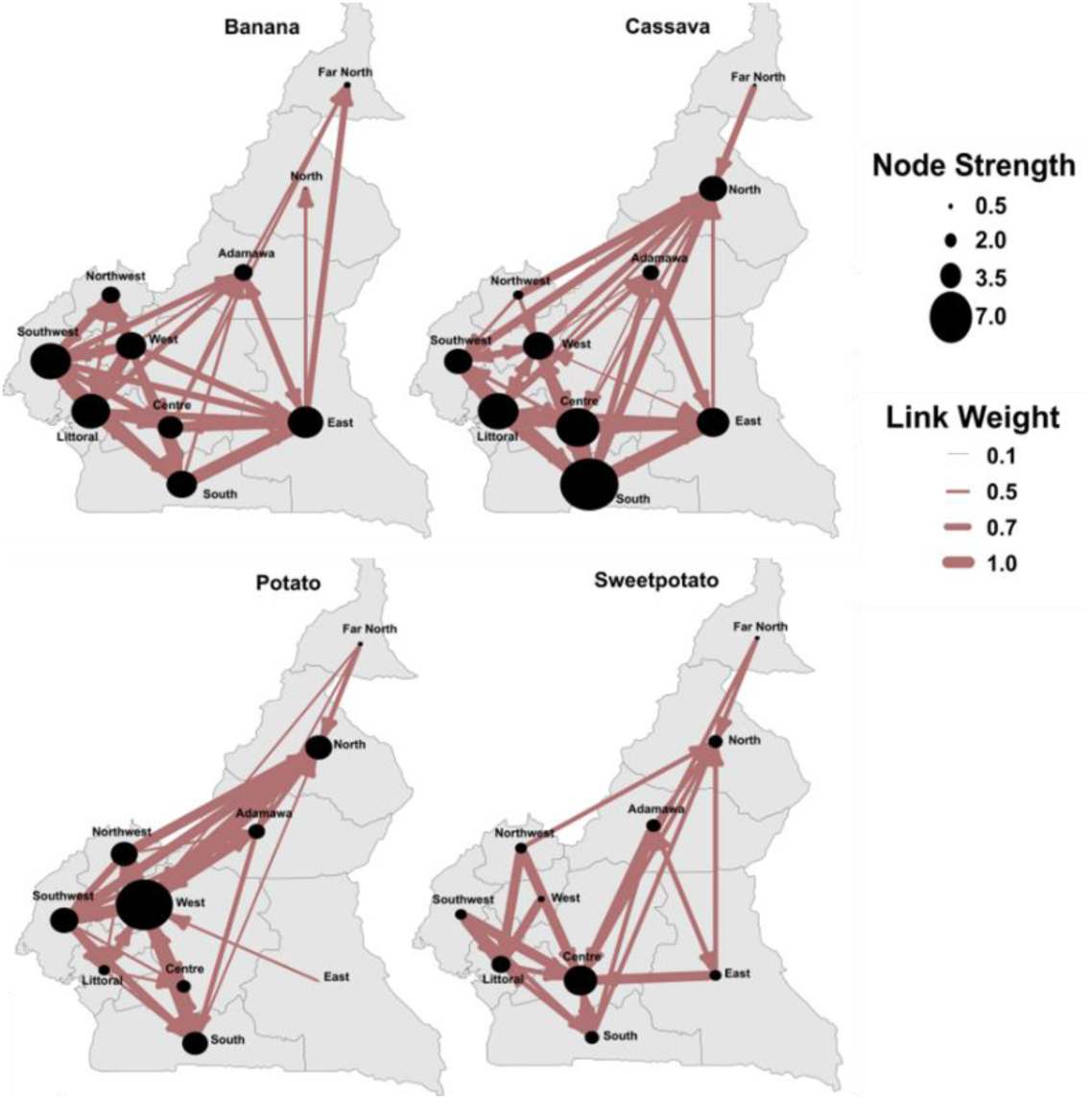
Informal seed trade estimated in expert knowledge elicitation suggests key regional hubs for potential within-country spread in Cameroon. Networks show reported informal planting-material exchange among regions for banana, cassava, potato, and sweetpotato. Node size indicates node strength (sum of link weights). Link weights represent the proportion of experts reporting informal trade between each pair of regions (i.e., confidence in the link), not trade volume or frequency. In post hoc discussions, some experts thought that detailed knowledge about seed flow and trade networks was not common among experts, so the uncertainty of these estimates may be higher. Movement of high quantities of perishable seed such as sweetpotato vines is likely to be uncommon at large distances.

For banana and cassava in Cameroon, regions were reported to be connected. Experts noted high regional informal seed trade strength for banana in the Southwest, Littoral, East, West and South regions for cassava in the South, Centre, Littoral, West and East regions (**Figure 4**). The experts also reported that, due to the current crisis in the Northwest and Southwest regions of Cameroon and the conflict with Boko Haram in the Far-North region, growers have moved to other regions and neighboring countries with their goods, including food and/or seed. Experts also highlighted the vulnerable situation of many of these growers working in neighboring regions and countries.

### 3.4 Informal seed trade with neighboring countries

We used expert knowledge elicitation to characterize informal seed trade networks that link Cameroon and Ethiopia with their neighboring countries. These networks were evaluated for potato and sweetpotato for both Cameroon and Ethiopia, and also for banana and cassava for Cameroon.

For informal banana seed in Cameroon, 40% of experts (4) noted exports to Gabon, and 30% (3) noted imports from Gabon and exports to Nigeria (**Figure 5, I**). Cameroon was noted to receive imports from Nigeria, the Central African Republic, Gabon, and the Republic of Congo, and noted to informally export to Chad, Nigeria, Central African Republic, Equatorial Guinea, Gabon, and the Republic of Congo.

**Figure 5.**
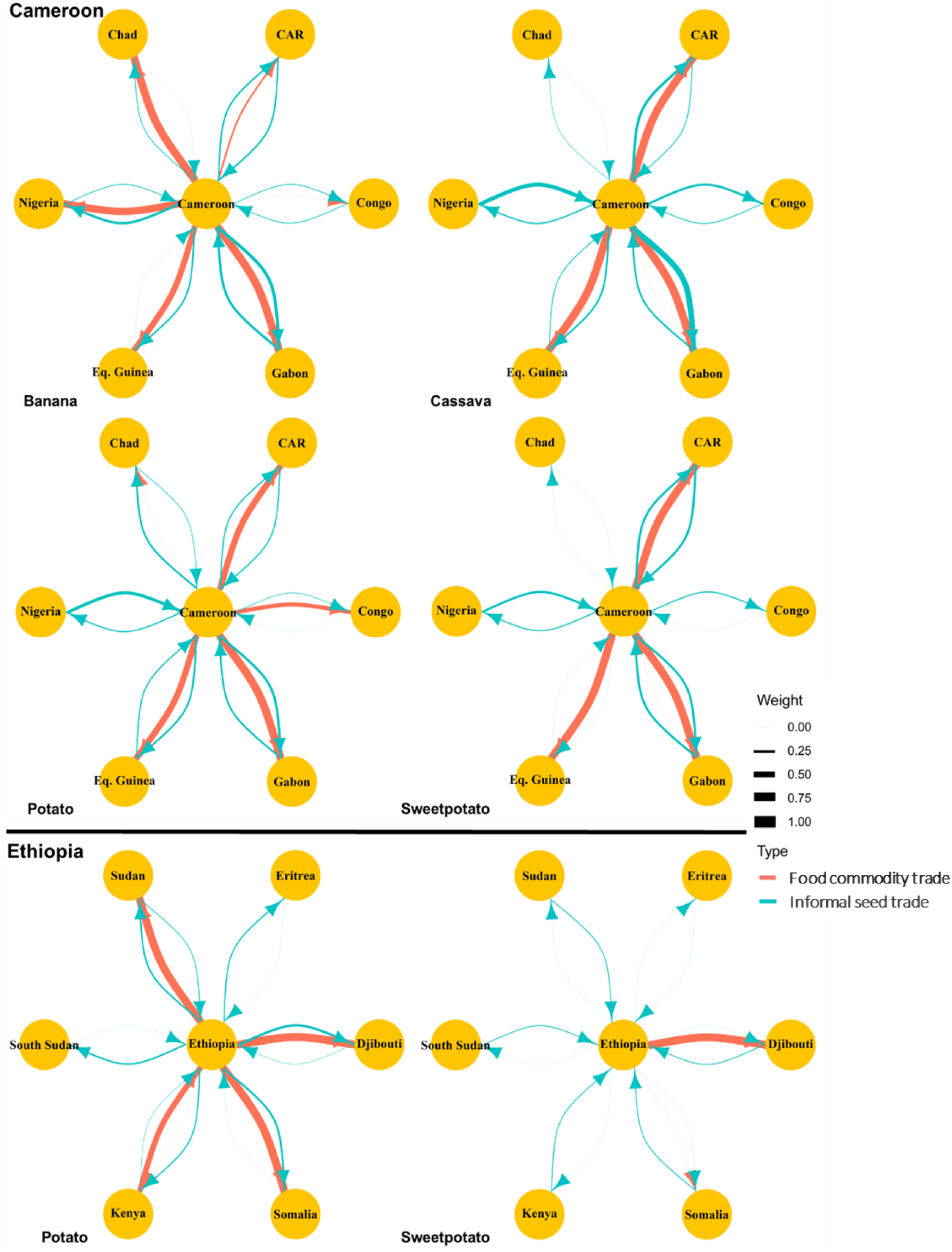
Cross-border networks highlight potential pathways for pathogen and pest movement via informal seed and formal commodity trade. Blue links show expert-reported informal seed trade with neighboring countries (link width proportional to the proportion of experts reporting each link). Red links show formal food commodity trade from FAO data (2006–2020 average; width proportional to trade volume). Food commodity trade may pose a particular risk for potato because ware tubers can be replanted as seed. CAR = Central African Republic, Congo = Republic of Congo, and Eq. Guinea = Equatorial Guinea.

For informal cassava seed trade in Cameroon, 72% of experts (13) noted exports to Gabon, and 50% (9) noted imports from Nigeria (**Figure 5, I**). Informal imports and exports were noted by experts for all neighboring countries, with the exception of informal imports from Chad (**Figure 5, I**).

For informal potato seed trade in Cameroon, 38% of experts (5) noted informal imports from Nigeria to Cameroon, and 30% (4) noted informal export from Cameroon to Gabon (**Figure 5, I**). Informal exports and imports of potato seed between all neighboring countries and Cameroon was noted by at least one expert, with the exception of imports from the Republic of Congo.

For informal sweetpotato seed imports, 29% of experts (2) noted that Gabon, Nigeria, and the Central African Republic imported from Cameroon, and 29% of experts (2) noted that Cameroon exported to Gabon and the Central African Republic (**Figure 5, I**). Cameroon was noted to informally export to Nigeria and Republic of Congo by 14% (1) of experts. No informal imports were noted from Chad, Equatorial Guinea, or the Republic of Congo, and no informal exports were noted to occur to Equatorial Guinea or Chad.

For informal Ethiopian potato seed trade, 30% of experts (6) noted export to Djibouti, and 25% (5) noted exports to Somalia and Sudan (**Figure 5, II**). No experts noted informal imports from Somalia, Eritrea, and South Sudan. Informal exports were noted for all neighboring countries. Informal trade of sweetpotato seed was reportedly rare from Ethiopia, where no informal exports of sweetpotato seed were noted (**Figure 5, II**). Informal imports were noted from all neighboring countries except Eritrea, with only 11% of experts (1) noting these imports.

### 3.5 Food commodity trade with neighboring countries

Food commodity trade is lower risk than informal seed trade, particularly for crops like banana and cassava where the food commodity cannot be used as planting material, but it still requires attention to ensure phytosanitary procedures. The potential for use of ware potatoes for planting is a particular risk. We used FAO trade data to characterize food commodity trade networks that link Cameroon and Ethiopia with their neighboring countries.

The highest volume trade for Cameroon in banana fruit was exports to Gabon, and secondly to Chad. Cameroon exported to Gabon, Chad, Nigeria, Equatorial Guinea, and the Central African Republic (**Figure 5, I**). For trade of cassava tubers in Cameroon, FAO reported the highest trade volume occurring as exports to Gabon. Exports were also to Equatorial Guinea and the Central African Republic (**Figure 5, I**). For Cameroonian trade in ware potato, only exports were reported with the Republic of Congo, the Central African Republic, Equatorial Guinea, and Gabon, with the highest trade volume average as exports to Gabon. For the trade in sweetpotato roots for consumption, bidirectional trade occurred between Cameroon and Gabon, with the highest trade volume occurred from Cameroon exports to Gabon. Cameroon also exports to the Central African Republic and Equatorial Guinea. No trade was reported to occur with Chad and the Republic of Congo (**Figure 5, I**). However, in follow up discussions, an expert in Cameroon reported observing large volumes of roots shipped from Cameroon to Chad daily, an illustration of how expert knowledge can supplement FAO reports.

For Ethiopian trade in ware potato, the highest trade volume was Ethiopian exports to Somalia, and then imports from Kenya. No imports or exports were reported between South Sudan and Eritrea, and no imports from Sudan, Djibouti, or Somalia (**Figure 5, II**). Sweetpotato tuber imports and exports were also reportedly rare, with Ethiopia exporting to Djibouti and Somalia (**Figure 5, II**).

## 4 Discussion

### 4.1 Components of this baseline national plant health risk analysis

Candidate priority locations, for surveillance and mitigation to reduce pathogen and pest risk, were identified based on cropland connectivity (**Figure 2**), informal seed trade (**Figures 4** and **5**), and commodity trade (**Figure 5**). Expert knowledge elicitation provided a geographic summary of major RTB pathogens and pests noted as present in the countries (**Figure 3**). These components of the national plant health risk analysis are intended to inform humanitarian and development actions, and to be improved over time as greater coordination is achieved between plant pathology research outputs and decision-making by humanitarian and development organizations. The baseline characterization of informal trade networks is also valuable for detecting how seed movement patterns shift during and after new crises, enabling more targeted responses.

The concept of technology readiness levels (TRLs) is helpful when considering the utility of this analysis for plant health management and perhaps especially humanitarian practitioners. The concept originated in aerospace development (Mankins, 2009), addressing the steps in development of a technology from TRL 1 (basic principles observed) to TRL 9 (actual system “flight proven” through successful operations). TRLs have since been adapted to evaluate technologies in crop development and disease diagnostics (USDA NIFA, 2018; White et al., 2022), and scaling readiness in research for development (Sartas et al., 2020). In humanitarian settings, operational context further modifies effective readiness: a technology proven at TRL 9 in a well-resourced setting may function at a much lower effective readiness in a disaster zone due to limited connectivity, power, infrastructure, and supply chains. Human readiness levels (HRLs), originally developed to complement TRLs, assess whether a technology is usable by its intended operators with available training (Phillips, 2010), a consideration directly relevant when tools must be usable by aid workers and affected communities with minimal support.

The analysis we present here is at approximately TRL 3, proof of concept. We demonstrate the potential to implement a rapid risk assessment as input for plant health management decision making, including in humanitarian projects. The current results may inform basic choices by humanitarian practitioners, such as selecting among equally priced seed source locations based on disease levels reported here. Although the present TRL is not sufficient to support decisions involving significantly differential costs or other trade-offs, we provide some examples of interpretation below.

Based on this TRL 3 analysis, cassava and sweetpotato trade between Cameroon and Nigeria warrants monitoring given high CCRI scores in both countries, while Ethiopia’s neighbors had low CCRI scores, suggesting lower cross-border risk from cropland connectivity alone (Buddenhagen et al., 2022; Saura et al., 2011). More direct information about trade fills in other dimensions of the potential for pathogen and pest movement (**Figures 4** and **5**). Given the extensive informal seed movement documented here—within regions, across regions, and across national borders—and the absence of phytosanitary controls, it can be assumed that informally traded planting material could carry pathogens and pests. The humanitarian context shapes both data and interpretation. Pulse (conflict, flooding) and press (instability, epidemics) stressors disrupt agricultural systems (Etherton et al., 2024). Ongoing crises in parts of Cameroon and Ethiopia mean reported cropland, expert perceptions, and trade patterns reflect disruption, not stable baselines. Under stability, cropland may be wider, trade routes more established, and knowledge more current; crises can also create new seed-movement pathways for pathogen and pest spread. Risk profiles therefore shift with changing conditions. Moreover, the experts’ cumulative years of experience in these countries span periods that include multiple crises, so their assessments already partially capture crisis-affected conditions rather than reflecting only stable circumstances. This makes the baseline useful both as a snapshot of partially disrupted systems and as a reference point against which future changes can be measured.

The current distribution of pathogens and pests is key for identifying crop biotic limitations and for understanding their potential spread to locations where they are currently absent or rare (**Figure 3**). For example, experts reported few pests and pathogens affecting banana, potato, and sweetpotato in East Cameroon, but all evaluated pathogens and pests were reported present in cassava. Some pathogens and pests such as white grub in Cameroon, were only reported in Centre and South Cameroon, and nowhere else in the country. Expert knowledge elicitation has been used in other systems to evaluate phytosanitary risk, for example by Andersen Onofre et al. (2021) to identify phytosanitary strategies for protecting potato seed systems in the Republic of Georgia. In a study linked to the one reported here, Etherton et al. (2025) created risk maps for potato seed systems in Ethiopia and evaluated regions at a high risk for pests and diseases, particularly *R. solanacearum*. Expert knowledge elicitation was also used to evaluate the informal trade network for seeds, and the risk of pathogen and pest spread between regions. Synthesizing across these analyses, locations where high cropland connectivity coincides with high reported pathogen and pest presence and active informal trade deserve particular attention. For example, in Cameroon, the West and Northwest regions show high CCRI scores for multiple crops (**Figure 2**), high diversity of reported pathogens and pests (**Figure 3**), and strong informal seed trade connections (**Figure 4**), making them candidate priorities for integrated surveillance and clean seed interventions. In Ethiopia, Oromia and Amhara similarly combine high potato cropland connectivity, diverse reported pathogen communities, and active trade connections. Future improvements to this analysis could formalize this synthesis through overlay analysis to generate composite risk maps, as in Etherton et al. (2025).

### 4.2 Ongoing improvement of a national plant health risk analysis

When resources are available, there are many ways to improve this national plant health risk analysis (**Figure 6**). For example, the cropland connectivity analyses represent a simplified view of the risk of spread by averaging across potential dispersal parameters, to represent many potential pathogens and pests. This broad parameterization prioritizes generality across many hazards, but at the cost of lower confidence for any single pathogen- or pest-specific dispersal scenario. When dispersal is well-characterized for a particular pathogen or pest species, the cropland connectivity analysis can be implemented specifically for that species, at the resolution needed, and potentially expanded to a fuller habitat connectivity analysis using the geohabnet package (Plex Sulá et al., 2025), including other geographic traits such as environmental conduciveness to the species. The cropland connectivity analysis may also be improved as crop density maps improve in quality (IFPRI, 2024; Tang et al., 2024). Cropland connectivity analyses can also be a first step for scenario analyses for evaluating regional pest and disease management strategies, by providing an estimate of the potential epidemic or invasion network for simulation experiments in scenario analyses (Etherton et al., 2025; Garrett, 2021; Wiebe et al., 2018). An important next step is to develop composite risk maps that overlay cropland connectivity, reported pathogen and pest distributions, and trade network intensity to identify locations where multiple risk factors converge, enabling more specific and testable predictions about disease spread risk.

**Figure 6.**
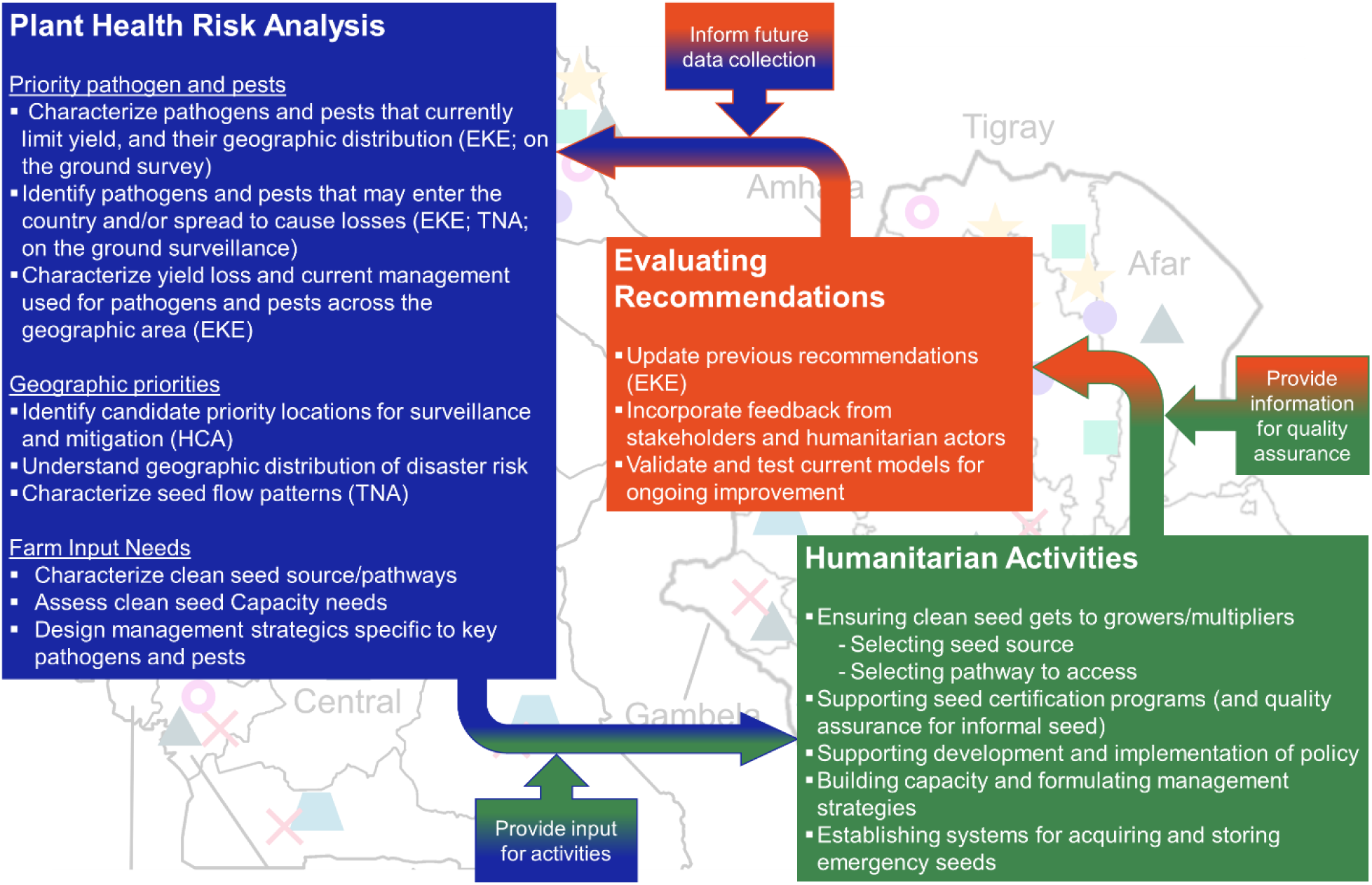
National plant health risk analysis framework to guide humanitarian seed and crop-health actions. The framework links plant health risk analysis (expert knowledge elicitation, habitat/cropland connectivity analysis, and trade network analysis) to humanitarian activities, with feedback loops to update recommendations and prioritize future data collection. EKE: expert knowledge elicitation, HCA: habitat/cropland connectivity analysis, and TNA: trade network analysis.

Expert knowledge elicitation can provide a baseline summary of expert perceptions, for ongoing improvement as new information becomes available. However, experts have gaps in their knowledge, leading to incomplete or biased information. There are also important cautionary aspects when expert knowledge elicitation is applied in policy contexts (Morgan, 2014). Experts may tend to have less knowledge about locations where a particular crop is less commonly produced, so that there is less ‘experiential sampling effort’ in those locations, in terms of expert knowledge. When resources are available for monitoring of pathogens and pests or trade, new information can supplement summaries of expert knowledge. Expert opinion can also serve as prior information in Bayesian analyses, updated as monitoring data accumulate (Ford et al., 1998; O’Hagan, 2019; Garthwaite et al., 2005; Hartley and French, 2021; Martin et al., 2005; Mkondiwa et al. 2024; Robledo et al., 2025; van de Schoot et al., 2021). In this study, we incorporate link weights indicating the relative confidence level across experts. After experts consider responses individually, discussion in small groups, as in methods like the Sheffield protocol for expert knowledge elicitation, can help refine expert opinions (Gosling, 2018; O’Hagan, 2019; Robledo et al., 2025). Future iterations could formalize monitoring priorities by combining expert-elicited uncertainty with value-of-information concepts to identify where new data would most reduce decision-relevant uncertainty (Runge et al., 2011; Buddenhagen et al., 2022). In practice, the appropriate level of investment in reducing uncertainty depends on the magnitude and cost of the decisions the analysis is intended to support (Morgan & Henrion, 1990).

Points for improvement in the formulation of questions for experts became apparent in this study. First, greater detail about the prevalence of pathogens and pests would be useful, in addition to presence/absence data, along with consideration of what experts anticipate could be emerging pathogens and pests in the near future. For example, future analyses should explicitly address high-priority emerging threats such as *Fusarium oxysporum* f.sp. *cubense* Tropical Race 4 (TR4) for banana. Second, the baseline information reported here addresses whether seed movement between locations occurs. Future evaluations could build on this by clarifying how frequent long-distance movement of perishable vegetative planting material is. The level of long-distance trade within a country is an important question from an epidemiological standpoint: long-distance pathogen movement can play a particularly important role in pathogen invasions across diverse regions. In follow up discussions, some experts thought that information about seed flow and trade networks was not common among crop and seed experts, so that there may be higher uncertainty associated with seed trade networks estimated using expert knowledge elicitation. Targeted follow-up studies addressing the quantity of seed moved through networks would be needed to decrease uncertainty. A third improvement concerns how disease importance is framed. Experts identified pathogens and pests as major problems, but the criteria for judging a disease a “problem”, and the severity or prevalence thresholds involved, were implicit. Making these criteria explicit in future elicitations would improve interpretability and comparability. We are documenting lessons on question formulation in annotated GitHub materials to guide future projects, and these lessons inform PlantQuest, the curated question catalog in the MetaQuestion web app (Fontan et al., 2025).

A national plant health risk analysis could also benefit from additional information and from links to larger and more general studies. For example, yield loss is an important component of risk, but detailed yield loss estimates are expensive. Savary et al. (2019) used an expert survey to estimate global yield losses due to pathogens and pests for five major food security crops, including potato. Information about the geographic distribution of current yield loss to pathogens and pest, and of management strategies adopted, is another potentially valuable component; these are also candidates for expert knowledge elicitation because experts with field experience often have useful knowledge but direct field surveys across large geographic areas are expensive to implement, especially when there are security issues for researchers (e.g., Mouafo-Tchinda et al., 2026). Expert knowledge elicitation and habitat connectivity analysis are part of the R2M Plant Health Toolbox for rapid risk assessment to support mitigation of plant pathogens and pests (garrettlab.com/r2m), designed for continual improvement based on lessons learned in applications across countries and crop systems. Future applications of expert knowledge elicitation can benefit from software supporting implementation (Hart et al., 2019). To facilitate projects addressing plant health, a new web application has been developed to provide a set of curated questions for expert knowledge elicitation related to plant health (Fontan et al., 2025). This tool, the R2M MetaQuestion web app drawing on the PlantQuest question catalog (Fontan et al., 2025), standardizes expert elicitation with curated questions about plant health and applied plant ecology, with a user interface that organizers can customize, including geographic pathogen/pest prevalence and planting material network analyses. These analyses can be used alongside related seed system tools (Sperling et al., 2022; Andrade-Piedra et al., 2023) and integrated into global initiatives such as the Global Plant Health Assessment (Acuña et al., 2023) and the proposed Global Surveillance System for Plant Disease (Carvajal-Yepes et al., 2019), and databases such as the CABI digital library (https://www.cabidigitallibrary.org/). Improvements can continually be made, in terms of the overall quality of data and modeling, and also in terms of the quality of translation for use in humanitarian applications.

### 4.3 Translating a national plant health risk analysis to support humanitarian activities

For RTB crops, a key point is supporting growers in obtaining quality seed, including growers displaced from neighboring regions or countries (Chapagain & Raizada, 2017; Etherton et al., 2024). Seed should have low pathogen and pest levels, and no pathogens or pests that would be new introductions to a location, to break or at least reduce cycles of seed degeneration (Moumni et al., 2023; Thomas-Sharma et al., 2017). Well-intentioned seed distribution can inadvertently introduce novel pests or pathogens into recipient areas if sourcing and phytosanitary measures are insufficient (McQuaid et al., 2016; Robson et al., 2024). Clean or certified seed is an expensive investment, and Cameroon and Ethiopia lack well-established system for distributing clean seed throughout the country. In Cameroon, for instance, 90% of the agricultural market, in general, is informal (OCHA, 2023a). Humanitarian practitioners should use caution in sourcing seed, drawing on information about the current distribution of pathogens and pests to reduce the chances of their introduction (**Figure 6**). It should be noted that because informal seed movement already occurs at scale and routinely carries pathogens and pests, humanitarian sourcing through the same channels may not substantially increase the marginal risk of pathogen spread beyond what is already occurring. The primary additional concern is the introduction of pathogens and pests that are new to a recipient area. Risk assessment of potential seed sources, initial seed testing (when stocks are obtained), maintenance of seed quality at multiple stages of the delivery chain, and charting of specific sources of original clean planting materials are all essential steps for humanitarian programs before and during on-the-ground interventions. Given these risks, disaster recovery with RTB crops should prioritize verifiable clean seed from formal seed sector channels where possible, a standard increasingly required by donors funding humanitarian agricultural interventions. Achieving a reliable supply of clean RTB planting material requires sustained investment in a seed value chain encompassing basic seed production, certified seed multiplication, and farmer-level seed management, a process requiring a minimum of three years to establish. In practice, the following two-step decision process can guide humanitarian seed responses for RTB crops.

The first step is to identify the risk of pathogen and pest movement via seed aid, using assessments such as those presented here. The second step is to decide whether to focus the response on the existing informal seed sector or to invest in establishing a formal seed value chain to ensure distributed seed is clean and free of pests and diseases. If the decision is to source seed from the informal sector, placing affected farmers at the center of the process (for example through cash voucher programs) can leverage their knowledge of acceptable seed quality and trusted local sources, and can be managed by humanitarian practitioners as a short-term activity. If instead the decision is to invest in a formal seed value chain, this requires greater resources per farmer reached, RTB-specific technical expertise, and a minimum of three years, and is best led by organizations with established RTB experience.

Because this analysis is at an early TRL, the resulting risk estimates should be treated as preliminary guidance rather than definitive assessments (Balafoutis et al., 2020). Humanitarian practitioners should weigh the confidence in these assessments against the consequences and costs of the decisions they inform (Nazemi et al., 2021). As the TRL advances through improved data and validation, confidence in such recommendations will increase and can more effectively support humanitarian-related decisions.

Providing clean seed to growers in locations with common seedborne pathogens and pests may be a candidate priority (Buddenhagen et al., 2022). Knowledge of seed systems and landscape connectivity can also be used to identify where the introduction of clean seed would produce the greatest benefit when pathogens or pests are actively invading (Andersen Onofre et al., 2025). In Ethiopia, candidate priority regions for potato clean seed aid include Oromia, SNNPR, Southwestern, Amhara, and Tigray (**Figure 3**). Cropland connectivity analysis, or more general habitat connectivity analysis, could be used before, during, and after crises to identify candidate priority locations where preventive or mitigation efforts are likely to have a greater impact on slowing pathogen or pest spread (Andersen Onofre et al., 2021; Thomas-Sharma et al., 2017; Xing et al., 2020). Maps of connectivity (CCRI) show surrounding regions that may also play an important role in epidemics, which may be priorities for providing clean seed to fleeing and displaced populations who are planting in new locations, or for seed testing to mitigate disease establishment. Similarly, locations that have informal trade with many other regions are also candidate priorities, to reduce dispersal of pathogens and pests, as, for example, the Southeast, South, West, and Centre regions of Cameroon for banana, cassava, potato, and sweetpotato, respectively (**Figure 4**). Following a crisis, understanding the informal movement of seed (**Figures 4** and **5**), as well as trade in food, can help identify management intervention priorities along informal trade chains, including growers, traders, and seed producers (Delaquis et al., 2018; Nduwimana et al., 2022).

A national crop health risk assessment can also support humanitarian activities beyond improvements to seed systems and seed sourcing. Disasters can influence pest and disease risk, and the potential for pathogen spread. For example, in Ethiopia, flooding in central Amhara (**Figure 1**) can occur along with conflicts such as the Tigray Civil conflict affecting the Amhara-Tigray border. Having an understanding of the pest and pathogen communities present in Amhara (**Figure 3**) can inform crop health priorities; when flooding occurs, aid could prioritize management of diseases like early blight, bacterial wilt, and late blight, which are exacerbated during floods (Etherton et al., 2024x; Guchi, 2015; Meno et al., 2020; Miley, 2020).

Capacity building can draw on information about important pathogens and pests in a priority region, and their potential movement. Training in the humanitarian practice community can address best practices for detecting and managing pathogens and pests, so that practitioners are prepared to evaluate seed quality and to provide training to farmers. Humanitarian practitioners often lack RTB-specific expertise and therefore need to partner with national and international RTB specialists to assess local seed system conditions, design appropriate interventions, and support implementation. Training could draw on the information in a national plant health risk analysis, and practitioners could provide feedback to improve the analysis and its utility to humanitarian personnel (**Figure 6**). If humanitarian practitioners can provide displaced farming families with basic information about identification of important pathogens and pests and their management, this can reduce the risk of spread to other farms and help to maintain yield.

Seed degeneration is likely exacerbated in Cameroon and Ethiopia since significant numbers of refugees and displaced people have been integrated into local communities, adding a further challenge to accessing clean seed and food. During crises, growers may abandon fields or flee, collapsing community pest and disease management. In stable periods, growers may collectively manage pests and pathogens at landscape scale. When this coordination breaks down, buildup accelerates and control becomes harder. There is often a lack of networks for accessing formal services and resources (ECPHAO, 2023; TGHAT, 2022).

Farm-to-farm transfer of planting material usually occurs at the village level, where recipient farmers could have first-hand knowledge of the health of their neighbors’ planting material (Cromwell, 1990; Sperling & Almekinders, 2023). Some of these sellers do have their own criteria for screening seed quality, like visual cleanliness of the planting material or its vigor, or its freshness of color. However, for many, screening for disease, especially diseases with more cryptic symptoms, is a formidable challenge. Farmers, traders, and humanitarian aid groups should be trained in selection for good-quality seed, combined with a strategy of regular seed replacement, which could be beneficial in areas with no formal seed system. This should be the first line of defense for safeguarding seed systems: education and training. More broadly, this study underscores the need for sustained collaboration between plant pathologists and humanitarian practitioners to develop practical, science-based solutions for agricultural resilience in crisis-affected regions (Rose & Adler, 2024).

## 4.4 Conclusions

Integrating the best available national plant health risk analysis in routine disaster risk reduction activities in countries experiencing multiple disasters can provide a baseline for rapid synthesis and verification during a crisis, directing practical humanitarian and government action. For on-farm management, the best available national plant health risk analysis can be used by extension specialists to design specific education and testing protocols in response to a disaster. At a larger scale, a national plant health risk analysis can be used by government agencies and humanitarian groups to allocate clean seed and management efforts both prior to and in response to disasters. This represents a novel effort and establishing a robust plant health risk analysis capacity in Cameroon, Ethiopia, and across Africa will require significant and reliable resources for a NARS-led effort, building on the baseline presented here. A national plant health risk analysis could also support long-term development efforts in peaceful periods, including targeted breeding for disease and pest resistance. This proactive approach could help vulnerable regions or countries to be better prepared for disasters and to recover from them more quickly. Aid, whether governmental or humanitarian, is a crucial part of agricultural sustainability and should focus on the development of sustainable long-term strategies, involving governmental agencies to develop more formal seed testing programs nationally. It is through the collaboration of humanitarian groups, governmental programs, national agricultural research institutions, traders, and farmers, themselves, that sustainable agriculture can be achieved in the face of humanitarian disasters.

## Statements and Declarations

The authors declare that they have no conflict of interest.

## Acknowledgements

We appreciate support from USAID Bureau for Humanitarian Assistance award number BHA 720BHA22IO00136 and the CGIAR Seed Equal Research Initiative; we thank all donors and organizations which globally support its work through their contributions to the CGIAR Trust Fund. We also appreciate support for the development of general methods from USDA Animal and Plant Health Inspection Service (APHIS) Cooperative Agreements AP21PPQS&T00C195 and AP22PPQS&T00C133. The opinions expressed in this article are those of the authors and do not necessarily reflect the view of the USAID BHA or USDA APHIS. We appreciate the contributions of participants in the expert knowledge elicitation, including experts who chose to remain anonymous and Gashaw Belay Alemu, Getachew Asefa, Atnafua Bekele, Tesfaye Abebe Desta, Astride S. Djabou Mouafi, Levai Lewis Dopgima, Demis Fikre, Dereje Haile, Wakene Hambisa, Denberu Kebede, Idriss Kouam, Bililign Mekonnen, Esmelealem Mihretu, Lionnel Nkamagne, Atsede Solomon, Yitagesu Tadesse, Lemlem Teklemedhin, Lemme Tessema, and Manamno Workayehu. We appreciate manuscript reviewers whose comments led to improvements in the manuscript.

## Reference

Acuña, I., Andrade-Piedra, J., Andrivon, D., Armengol, J., Elizabeth Arnold, A., Avelino, J., et al. (2023). A global assessment of the state of plant health. Plant Disease, 107(12), 3649–3665.

Alasow, A. A., Hamed, M. M., & Shahid, S. (2023). Spatiotemporal variability of drought and affected croplands in the Horn of Africa. Stochastic Environmental Research and Risk Assessment, 1–16. 10.1007/s00477-023-02575-1

Andersen, K. F., Buddenhagen, C. E., Rachkara, P., Gibson, R., Kalule, S., Phillips, D., and Garrett, K. A. (2019). Modeling epidemics in seed systems and landscapes to guide management strategies: The case of sweetpotato in Northern Uganda. Phytopathology, 109, 1519–1532.

Andersen Onofre, K. F., Delaquis, E., Newby, J., de Haan, S., Thuy, C. T. L., Minato, N., Legg, J., Cuellar, W., Alcalá Briseño, R. I., & Garrett, K. A. (2025). Decision support for managing an invasive pathogen through efficient clean seed systems: Cassava mosaic disease in Southeast Asia. Agricultural Systems, 229, 104435.

Andersen Onofre, K. F., Forbes, G. A., Andrade-Piedra, J. L., Buddenhagen, C. E., Fulton, J. C., Gatto, M., Khidesheli, Z., Mdivani, R., Xing, Y., & Garrett, K. A. (2021). An integrated seed health strategy and phytosanitary risk assessment: Potato in the Republic of Georgia. Agricultural Systems, 191, 103144. 10.1016/j.agsy.2021.103144

Andrade-Piedra, J. L., Sharma, K., Kroschel, J., Ogero, K., Kreuze, J., Legg, J. P., Kumar, P. L., Spielman, D. J., Navarrete, I., Perez, W., Atieno, E., & Garrett, K. A. (2025). Phytosanitary challenges and solutions for roots and tubers in the tropics. Annual Review of Phytopathology, 63, 627–650.

Andrade-Piedra, J., Dontsop, P., Dunia, D., Kulakow, P., Harahagazwe, D., Mouafo-Tchinda, R., Omondi, A., Ogero, K., Rajendran, S., & McEwan, M. (2023). Tools4SeedSystems: Working towards resilience through root, tuber and banana crops in humanitarian settings. Capacity needs assessment for root, tuber and banana seed interventions in humanitarian settings: Cameroon and DRC. Technical Report. 10.4160/9789290606635

Arndt, E., Rumpff, L., Lane, S., Bau, S., Mebalds, M., and Kompas, T. (2022). Estimating probability of visual detection of exotic pests and diseases in the grains industry—An expert elicitation approach. Front. Ecol. Evol., 10, 968436 10.3389/fevo.2022.968436.

Balafoutis, A. T., Evert, F. K. V., & Fountas, S. (2020). Smart farming technology trends: Economic and environmental effects, labor impact, and adoption readiness. Agronomy, 10(5), 743. 10.3390/agronomy10050743

Bang, H. N. (2021). A gap analysis of the legislative, policy, institutional and crises management frameworks for disaster risk management in Cameroon. Progress in Disaster Science, 111, 00190. 10.1016/j.pdisas.2021.100190

Bang, H. N., & Balgah, R. A. (2022). The ramification of Cameroon’s Anglophone crisis: conceptual analysis of a looming “Complex Disaster Emergency”. Journal of International Humanitarian Action, 7(1), 6.

Battiston, P., Kashyap, R., and Rotondi, V. (2021). Reliance on scientists and experts during an epidemic: Evidence from the COVID-19 outbreak in Italy. SSM - Population Health, 13, 100721. 10.1016/j.ssmph.2020.100721

Beyene, K. (2015). Destitution, biology, yield loss and management of sweet potato weevils (*Cylas formicaries* (Fabrcius) Insecta: Coleoptera) in Ethiopia. Journal of Biology, Agriculture and Healthcare, 5(22), 65–72.

Bramel, P., Nagoda, S., Haugen, J., Adugna, D., Dejene, T., Bekele, T., & Traedal, L. T. (2004). Relief seed assistance in Ethiopia. Pages 111–133 in Addressing Seed Security in Disaster Response: Linking Relief with Development. CIAT Cali.

Buddenhagen, C. E., Xing, Y., Andrade-Piedra, J. L., Forbes, G. A., Kromann, P., Navarrete, I., Thomas-Sharma, S., Choudhury, R. A., Andersen Onofre, K. F., Schulte-Geldermann, E., Etherton, B. A., Plex Sula, A. I., & Garrett, K. A. (2022). Where to invest project efforts for greater benefit: A framework for management performance mapping with examples for potato seed health. Phytopathology, 112(7), 1431–1443. 10.1094/PHYTO-05-20-0202-R

Canney-Davison, S., Davies, L., & McEwan, M. (2023). Tools4SeedSystems: Working towards resilience through root, tuber and banana crops in humanitarian settings. Workshop Report.

Carvajal-Yepes, M., Cardwell, K., Nelson, A., Garrett, K. A., Giovani, B., Saunders, D., Kamoun, S., Legg, J., Verdier, V., & Lessel, J. (2019). A global surveillance system for crop diseases. Science, 364(6447), 1237–1239. 10.1126/science.aaw1572

CDP. (2023). Ethiopia humanitarian crisis. https://disasterphilanthropy.org/disasters/ethiopia-tigray-crisis/

Chapagain, T., & Raizada, M. N. (2017). Impacts of natural disasters on smallholder farmers: gaps and recommendations. Agriculture & Food Security, 6, 1–16.

Chen, M., Brun, F., Raynal, M., Debord, C., and Makowski, D. (2019). Use of probabilistic expert elicitation for assessing risk of appearance of grape downy mildew. Crop Protection, 126, 104926. 10.1016/j.cropro.2019.104926

Colson, A. R., and Cooke, R. M. (2018). Expert elicitation: Using the classical model to validate experts’ judgments. Review of Environmental Economics and Policy, 12, 113–132. 10.1093/reep/rex022

Cooke, R. M. (1991). Experts in Uncertainty: Opinion and Subjective Probability in Science. New York, NY, US: Oxford University Press.

Cooke, R. M., and Goossens, L. H. J. (2004). Expert judgement elicitation for risk assessments of critical infrastructures. Journal of Risk Research, 7, 643–656. 10.1080/1366987042000192237

Cromwell, E. (1990). Seed diffusion mechanisms in small farmer communities: Lessons from Asia, Africa and Latin America.

Csárdi, G., Nepusz, T., Traag, V., Horvát, S., Zanini, F., Noom, D., & Müller, K. (2023). igraph: Network Analysis and Visualization in R. R package version 1.5.1.

Delaquis, E., Andersen, K. F., Minato, N., Cu, T. T. L., Karssenberg, M. E., Sok, S., Wyckhuys, K. A., Newby, J. C., et al. (2018). Raising the stakes: cassava seed networks at multiple scales in Cambodia and Vietnam. Frontiers in Sustainable Food Systems, 2, 73. 10.3389/fsufs.2018.00073

Dicko, A. (2023). rgeoboundaries: A Client to geoBoundaries, A Political Administrative Boundaries

Doungous, O., Masky, B., Levai, D. L., Bahoya, J. A., Minyaka, E., Mavoungou, J. F., Mutuku, J. M., & Pita, J. S. (2022). Cassava mosaic disease and its whitefly vector in Cameroon: Incidence, severity and whitefly numbers from field surveys. Crop Protection, 158, 106017. 10.1016/j.cropro.2022.106017

ECPHAO. (2023). Cameroon factsheet: European Civil Protection and Humanitarian Aid Operations. https://civil-protection-humanitarian-aid.ec.europa.eu/where/africa/cameroon_en

EFSA Panel on Plant Health, Jeger, M., Bragard, C., Caffier, D., Candresse, T., Chatzivassiliou, E., Dehnen-Schmutz, K., Grégoire, J., Jaques Miret, J. A., and MacLeod, A. (2018). Guidance on quantitative pest risk assessment. EFSA Journal, 16, e05350.

Etherton, B. A., Choudhury, R. A., Alcalá Briseño, R. I., Mouafo-Tchinda, R. A., Plex Sulá, A. I., Choudhury, M., Adhikari, A., Lei, S. L., Kraisitudomsook, N., & Buritica, J. R. (2024). Disaster plant pathology: Smart solutions for threats to global plant health from natural and human-driven disasters. Phytopathology, 114(5), 855–868.

Etherton, B., Plex Sula, A., Mouafo-Tchinda, R., Kakuhenzire, R., Kassaye, H., Asfaw, F., Kosmakos, V., McCoy, R., Xing, Y., & Yao, J. (2025). Translating Ethiopian potato seed networks: Identifying strategic intervention points for managing bacterial wilt and other diseases. Agricultural Systems, 222, 104167. 10.1016/j.agsy.2024.104167

European Food Safety Authority. 2014. Guidance on expert knowledge elicitation in food and feed safety risk assessment. EFSA Journal, 12, 3734.

FAO. (2021a). Emergency assistance to improve food and nutrition security of IDPS in South West region of Cameroon. https://www.fao.org/documents/card/fr/c/CB3577EN/

FAO. (2021b). The impact of disasters and crises on agriculture and food security. https://www.fao.org/documents/card/en/c/cb3673en

FAOSTAT. (2021). Trade: crops and livestock products. https://www.fao.org/faostat/en/#home

Fontan, R., Perez, C. M., Adhikari, A., Mouafo-Tchinda, R. A., Plex Sulá, A. I., Robledo, J., Etherton, B. A., Choudhary, M., Sarwar, M. A., Naveed, Z. A., & Garrett, K. A. (2025). MetaQuestion: A web application for expert knowledge elicitation addressing plant health and applied plant ecology. arXiv 2509.19393.

Ford, J. K., Smith, E. M., Weissbein, D. A., Gully, S. M., & Salas, E. (1998). Relationships of goal orientation, metacognitive activity, and practice strategies with learning outcomes and transfer. Journal of Applied Psychology, 83(2), 218.

Foyou, V. E., Ngwafu, P., Santoyo, M., & Ortiz, A. (2018). The Boko Haram insurgency and its impact on border security, trade and economic collaboration between Nigeria and Cameroon: An exploratory study. African Social Science Review, 9(1), 7. https://digitalscholarship.tsu.edu/assr/vol9/iss1/7

Garrett, K. A. (2021). Impact network analysis and the INA R package: Decision support for regional management interventions. Methods in Ecology and Evolution, 12, 1634–1647.

Garrett, K. A., Bebber, D., Etherton, B. A., Gold, K., Plex Sulá, A. I., & Selvaraj, M. G. (2022). Climate change effects on pathogen emergence: artificial intelligence to translate big data for mitigation. Annual Review of Phytopathology, 60, 357–378. 10.1146/annurev-phyto-021021-042636

Garthwaite, P. H., Kadane, J. B., and O’Hagan, A. (2005). Statistical methods for eliciting probability distributions. Journal of the American Statistical Association, 100, 680–701. 10.1198/016214505000000105

Gibson, R., & Kreuze, J. F. (2015). Degeneration in sweetpotato due to viruses, virus-cleaned planting material and reversion: a review. Plant Pathology, 64(1), 1–15. 10.1111/ppa.12273

Gilligan, C. A. (2008). Sustainable agriculture and plant diseases: An epidemiological perspective. Philosophical Transactions of the Royal Society B: Biological Sciences, 363(1492), 741–759.

Gosling, J. P. (2018). SHELF: the Sheffield elicitation framework. Elicitation: The science and art of structuring judgement, 61–93. 10.1007/978-3-319-65052-4_4

Gosling, J. P. (2018). SHELF: The Sheffield Elicitation Framework. In Elicitation: The Science and Art of Structuring Judgement, eds. L. C. Dias, A. Morton, and J. Quigley. Cham: Springer International Publishing, pp. 61–93. 10.1007/978-3-319-65052-4_4

Guchi, E. (2015). Disease management practice on potato (*Solanum tuberosum* L.) in Ethiopia. World Journal of Agricultural Research, 3(1), 34–42.

Hadjigeorgiou, E., Clark, B., Simpson, E., Coles, D., Comber, R., Fischer, A. R. H., Meijer, N., Marvin, H. J. P., and Frewer, L. J. (2022). A systematic review into expert knowledge elicitation methods for emerging food and feed risk identification. Food Control, 136, 108848. 10.1016/j.foodcont.2022.108848

Hart, C. M. P. ’t, Leontaris, G., and Morales-Nápoles, O. (2019). Update (1.1) to ANDURIL — A MATLAB toolbox for ANalysis and Decisions with UnceRtaInty: Learning from expert judgments: ANDURYL. SoftwareX, 10, 100295. 10.1016/j.softx.2019.100295

Hartley, D., and French, S. (2021). A Bayesian method for calibration and aggregation of expert judgement. International Journal of Approximate Reasoning, 130, 192–225. 10.1016/j.ijar.2020.12.007

Hemming, V., Burgman, M. A., Hanea, A. M., McBride, M. F., and Wintle, B. C. (2018). A practical guide to structured expert elicitation using the IDEA protocol. Methods in Ecology and Evolution, 9, 169–180. 10.1111/2041-210X.12857

Hirpa, A., Meuwissen, M. P., Tesfaye, A., Lommen, W. J., Lansink, A. O., Tsegaye, A., & Struik, P. C. (2010). Analysis of seed potato systems in Ethiopia. American Journal of Potato Research, 87(6), 537–552. 10.1007/s12230-010-9164-1

Hughes, G., and Madden, L. V. (2002). Some methods for eliciting expert knowledge of plant disease epidemics and their application in cluster sampling for disease incidence. Crop Protection, 21, 203–215. 10.1016/S0261-2194(01)00087-4

Hunt, M., Chénier, A., Bezanson, K., Nouvet, E., Bernard, C., de Laat, S., Krishnaraj, G., & Schwartz, L. (2018). Moral experiences of humanitarian health professionals caring for patients who are dying or likely to die in a humanitarian crisis. Journal of International Humanitarian Action, 3(1), 1–13. 10.1186/s41018-018-0040-9

IFPRI. (2020). Spatially-Disaggregated Crop Production Statistics Data in Africa South of the Sahara for 2017. Harvard Dataverse International Food Policy Research Institute.

IFPRI. (2020). Spatially-disaggregated crop production statistics data in Africa south of the Sahara for 2017, 10.7910/DVN/FSSKBW [Spatially-disaggregated crop production statistics]. Harvard Dataverse, V2. doi:10.7910/DVN/FSSKBW

IFPRI. (2024). Global Spatially-Disaggregated Crop Production Statistics Data for 2020 Version 1.0. In: Harvard Dataverse, V3.

IOM. (2023). More than 4.38 million people displaced in Ethiopia, more than half due to conflict: New IOM Report. https://ethiopia.iom.int/news/more-438-million-people-displaced-ethiopia-more-half-due-conflict-new-iom-report

Jeger, M. J., Pautasso, M., Holdenrieder, O., & Shaw, M. W. (2007). Modelling disease spread and control in networks: implications for plant sciences. New Phytologist, 174(2), 279–297.

Jeger, M., Holt, J., Van Den Bosch, F., & Madden, L. (2004). Epidemiology of insect-transmitted plant viruses: modelling disease dynamics and control interventions. Physiological Entomology, 29(3), 291–304.

Jongejans, E., Skarpaas, O., Ferrari, M. J., Long, E. S., Dauer, J. T., Schwarz, C. M., Rauschert, E. S., Jabbour, R., Mortensen, D. A., & Isard, S. A. (2015). A unifying gravity framework for dispersal. Theoretical Ecology, 8(2), 207–223. 10.1007/s12080-014-0245-5

Knol, A. B., Slottje, P., van der Sluijs, J. P., and Lebret, E. (2010). The use of expert elicitation in environmental health impact assessment: a seven step procedure. Environmental Health, 9, 19. 10.1186/1476-069X-9-19

Konje, C., Abdulai, A., Tange, A. D., Tarla, D., & Tita, M. A. (2019). Identification and management of pests and diseases of garden crops in Santa, Cameroon. Journal of Agriculture and Ecology Research International, 18(2), 1–9. 10.9734/JAERI/2019/v18i230055

Kwambai, T. K., Griffin, D., Struik, P. C., Stack, L., Rono, S., Brophy, C., Nyongesa, M., & Gorman, M. (2024). Seed quality and variety preferences amongst potato farmers in North-Western Kenya: lessons for the adoption of new varieties. Potato Research, 67(1), 185–208.

Macrotrends. (2023). Cameroon Refugee Statistics 1970-2023. https://www.macrotrends.net/countries/CMR/cameroon/refugee-statistics

Man, C. R. (2018). Seed aid, smallholders, and the developmental state: A mixed methods analysis of emergency relief programs in Ethiopia. The Pennsylvania State University.

Mankins, J. C. (2009). Technology readiness assessments: A retrospective. Acta Astronautica, 65, 1216–1223.

Margosian, M. L., Garrett, K. A., Hutchinson, J. S., & With, K. A. (2009). Connectivity of the American agricultural landscape: assessing the national risk of crop pest and disease spread. BioScience, 59(2), 141–151. 10.1525/bio.2009.59.2.7

Martin, T. G., Burgman, M. A., Fidler, F., Kuhnert, P. M., Low-Choy, S., Mcbride, M., and Mengersen, K. (2012). Eliciting expert knowledge in conservation science. Conservation Biology, 26, 29–38. 10.1111/j.1523-1739.2011.01806.x

Martin, T. G., Kuhnert, P. M., Mengersen, K., and Possingham, H. P. (2005). The power of expert opinion in ecological models using Bayesian methods: Impact of grazing on birds. Ecological Applications, 15, 266–280. 10.1890/03-5400

Mbah, L. T., Molua, E. L., Bomdzele, E., & Egwu, B. M. (2023). Farmers’ response to maize production risks in Cameroon: An application of the criticality risk matrix model. Heliyon, 9(4), e15124. 10.1016/j.heliyon.2023.e15124

McQuaid, C., Sseruwagi, P., Pariyo, A., and Van den Bosch, F. (2016). Cassava brown streak disease and the sustainability of a clean seed system. Plant Pathology, 65, 299–309.

Mengistu, D. (2001). Linking emergency aid with rehabilitation and support in chronic stress situations. Targeted Seed Aid and Seed-System Interventions, 59.

Meno, L., Escuredo, O., Rodríguez-Flores, M. S., & Seijo, M. C. (2020). Modification of the TOMCAST model with aerobiological data for management of potato early blight. Agronomy, 10(12), 1872.

Mila, A. L., and Carriquiry, A. L. (2004). Bayesian analysis in plant pathology. Phytopathology, 94, 1027–1030. 10.1094/PHYTO.2004.94.9.1027

Miley, K. (2020). Global health biosecurity in a vulnerable world–an evaluation of emerging threats and current disaster preparedness strategies for the future. Global Health Security: Recognizing Vulnerabilities, Creating Opportunities, 79–102.

Mkondiwa, M., Hurley, T. M., and Pardey, P. G. (2024). Closing the gaps in experimental and observational crop response estimates: a Bayesian approach. Q Open, 4, qoae017. 10.1093/qopen/qoae017

Morgan, M. G. (2014). Use (and abuse) of expert elicitation in support of decision making for public policy. Proceedings of the National Academy of Sciences, 111, 7176–7184. 10.1073/pnas.1319946111

Morgan, M. G., & Henrion, M. (1990). Uncertainty: A Guide to Dealing with Uncertainty in Quantitative Risk and Policy Analysis. Cambridge University Press.

Motisi, N., Bommel, P., Leclerc, G., Robin, M.-H., Aubertot, J.-N., Butron, A. A., Merle, I., Treminio, E., and Avelino, J. (2022). Improved forecasting of coffee leaf rust by qualitative modeling: Design and expert validation of the ExpeRoya model. Agricultural Systems, 197,103352. 10.1016/j.agsy.2021.103352

Mouafo-Tchinda, R. A., Plex Sulá, A. I., Etherton, B. A., Okonya, J. S., Nakato, G. V., Xing, Y., Jacobo, R. B., Adhikari, A., Blomme, G., Kantungeko, D., Nduwayezu, A., Kreuze, J., Kroschel, J., Legg, J., and Garrett, K. A. 2026. Pathogen and pest communities in food security crops across climate gradients: Anticipating future challenges in the highland tropics. Agricultural Systems, 233, 104619.

Moumni, M., Brodal, G., & Romanazzi, G. (2023). Recent innovative seed treatment methods in the management of seedborne pathogens. Food Security, 15(5), 1365–1382.

Nanbol, K. K., & Namo, O. (2019). The contribution of root and tuber crops to food security: A review. J. Agric. Sci. Technol. B, 9(10.17265), 2161–6264.

Nazemi, N., Parragh, S. N., & Gutjahr, W. J. (2020). Bi-objective facility location under uncertainty with an application in last-mile disaster relief. Annals of Operations Research, 319(2), 1689–1716. 10.1007/s10479-021-04422-4

Nduwimana, I., Sylla, S., Xing, Y., Simbare, A., Niyongere, C., Garrett, K. A., & Bonaventure Omondi, A. (2022). Banana seed exchange networks in Burundi–Linking formal and informal systems. Outlook on Agriculture, 51(3), 334–348. 10.1177/00307270221103288

Ngatat, S., Hanna, R., Doumtsop Fotio, A. R., Lienou, J. A., Nanga Nanga, S., Fotso Kuate, A., Osundahunsi, B., Fiaboe, K. K., Ndemba, B., & Dossa, G. S. (2024). Distribution and diversity of emergent Banana bunchy top virus infecting banana and plantain in Cameroon, Central Africa. Journal of Phytopathology, 172(2), e13279.

Nisbet, C., Lestrat, K. E., & Vatanparast, H. (2022). Food security interventions among refugees around the globe: a scoping review. Nutrients, 14(3), 522.

O’Hagan, A. (2019). Expert knowledge elicitation: subjective but scientific. The American Statistician, 73(sup1), 69–81. 10.1080/00031305.2018.1518265

OCHA. (2022). Cameroon multi-stakeholder dialogue on refugees.https://reliefweb.int/report/cameroon/cameroon-multi-stakeholder-dialogue-refugees-november-2022

OCHA. (2023a). Cameroon humanitarian needs overview 2023. https://www.unocha.org/publications/report/cameroon/cameroon-humanitarian-needs-overview-2023-march-2023

OCHA. (2023b). Ethiopia: humanitarian needs overview 2023. https://reliefweb.int/report/ethiopia/ethiopia-humanitarian-needs-overview-2023

Pedersen, T. L. (2022). ggraph: An Implementation of Grammar of Graphics for Graphs and Networks. Retrieved January, 1, 2018. https://CRAN.R-project.org/package=ggraph

Petsakos, A., Prager, S. D., Gonzalez, C. E., Gama, A. C., Sulser, T. B., Gbegbelegbe, S., Kikulwe, E. M., & Hareau, G. (2019). Understanding the consequences of changes in the production frontiers for roots, tubers and bananas. Global Food Security, 20, 180–188.

Plex Sulá, A. I., Keshav, K., Adhikari, A., Mouafo-Tchinda, R. A., Robledo, J., Shah, S. N., and Garrett, K. A. (2025). geohabnet: An R package for mapping habitat connectivity for biosecurity and conservation. arXiv 2510.24955.

Phillips, E. L. 2010. “The Development and Initial Evaluation of the Human Readiness Level Framework.” Naval Postgraduate School.

Poubom, C., Awah, E., Tchuanyo, M., & Tengoua, F. (2005). Farmers’ perceptions of cassava pests and indigenous control methods in Cameroon. International Journal of Pest Management, 51(2), 157–164. 10.1080/09670870500131863

Prasanna, B. M., Carvajal-Yepes, M., Kumar, P. L., Kawarazuka, N., Liu, Y., Mulema, A. A., McCutcheon, S., & Ibabao, X. (2022). Sustainable management of transboundary pests requires holistic and inclusive solutions. Food Security, 14(6), 1449–1457.

Raleigh, C., Linke, R., Hegre, H., & Karlsen, J. (2010). Introducing ACLED: An armed conflict location and event dataset. Journal of Peace Research, 47(5), 651–660. 10.1177/0022343310378914

Robledo, J., Plex Sulá, A. I., Jaworski, L. G., Mouafo-Tchinda, R. A., Andersen Onofre, K. F., Thomas-Sharma, S., and Garrett, K. A. (2025). Expert knowledge elicitation: Accessing the big data in experts’ brains. Phytopathology 115:1245–1259.

Robson, F., Hird, D. L., and Boa, E. (2024). Cassava brown streak: A deadly virus on the move. Plant Pathology, 73, 21–241.

Rose, J., & Adler, C. M. (2024). A framework for effective collaboration with crisis-affected communities. Challenges, 15(1), 13. 10.3390/challe15010013

Rose, L. E., Hemming, V., Hanea, A. M., Wintle, B. A., and Chee, Y. E. (2023). Linking species distribution models with structured expert elicitation for predicting management effectiveness. Conservation Science and Practice, 5, e13038. 10.1111/csp2.13038

Runge, M. C., Converse, S. J., and Lyons, J. E. (2011). Which uncertainty? Using expert elicitation and expected value of information to design an adaptive program. Biological Conservation, 144, 1214–1223. 10.1016/j.biocon.2010.12.020

Runfola, D., Anderson, A., Baier, H., Crittenden, M., Dowker, E., Fuhrig, S., et al. (2020). geoBoundaries: A global database of political administrative boundaries. PLoS ONE, 15(4), e0231866.

Sartas, M., Schut, M., Proietti, C., Thiele, G., and Leeuwis, C. (2020). Scaling readiness: science and practice of an approach to enhance impact of research for development. Agricultural Systems, 183, 102874.

Saura, S., Vogt, P., Velázquez, J., Hernando, A., & Tejera, R. (2011). Key structural forest connectors can be identified by combining landscape spatial pattern and network analyses. Forest Ecology and Management, 262(2), 150–160.

Savary, S., Willocquet, L., Pethybridge, S. J., Esker, P., McRoberts, N., & Nelson, A. (2019). The global burden of pathogens and pests on major food crops. Nature Ecology & Evolution, 3(3), 430–439. 10.1038/s41559-018-0793-y

Scott, G. J. (2021). A review of root, tuber and banana crops in developing countries: past, present and future. International Journal of Food Science & Technology, 56(3), 1093–1114. 10.1111/ijfs.14778

Shonga, E., Gemu, M., Tadesse, T., & Urage, E. (2013). Review of entomological research on sweet potato in Ethiopia. Discourse Journal of Agriculture and Food Sciences, 1(5), 83–92.

Singh, R. P., Prasad, P. V. V., & Reddy, K. R. (2015). Climate change: implications for stakeholders in genetic resources and seed sector. Advances in Agronomy, 129, 117–180.

Soares, M., Colson, A., Bojke, L., Ghabri, S., Garay, O. U., Felli, J. K., Lee, K., Molsen-David, E., Morales-Napoles, O., Shaffer, V. A., and Ijzerman, M. J. (2024). Recommendations on the use of structured expert elicitation protocols for healthcare decision making: A good practices report of an ISPOR task force. Value in Health, 27, 1469–1478. 10.1016/j.jval.2024.07.027

Sperling, L. (2008). When disaster strikes: a guide to assessing seed system security. CIAT.

Sperling, L., & McGuire, S. J. (2010). Persistent myths about emergency seed aid. Food Policy, 35(3), 195–201.

Sperling, L., & Almekinders, C. J. (2023). Informal commercial seed systems: Leave, suppress or support them? Sustainability, 15(18), 14008. 10.3390/su151814008

Sperling, L., Gallagher, P., McGuire, S., March, J., & Templer, N. (2020). Informal seed traders: the backbone of seed business and African smallholder seed supply. Sustainability, 12(17), 7074.

Sperling, L., Mottram, A., Ouko, W., & Love, A. (2022). Seed emergency response tool: Guidance for practitioners. Produced by Mercy Corps and SeedSystem. org as a Part of the ISSD Africa Activity.

Szyniszewska, A. M., Chikoti, P. C., Tembo, M., Mulenga, R., Gilligan, C. A., van den Bosch, F., and McQuaid, C. F. (2021). Smallholder cassava planting material movement and grower behavior in Zambia: implications for the management of cassava virus diseases. Phytopathology, 111, 1952–1962.

Tang, F. H., Nguyen, T. H., Conchedda, G., Casse, L., Tubiello, F. N., & Maggi, F. (2024). CROPGRIDS: a global geo-referenced dataset of 173 crops. Scientific Data, 11(1), 413.

Tessema, G. L., & Seid, H. E. (2023). Potato bacterial wilt in Ethiopia: history, current status, and future perspectives. PeerJ, 11, e14661. 10.7717/peerj.14661

Tessema, L., & Tesfaye, M. (2023). Understanding and managing seed degeneration in potato: Implications for potato resilient seed system and food security. A review. CABI Reviews.

TGHAT. (2022). Tigray bureau of agriculture and natural resources, emergency and recovery support plan for the 2022/23 budget year, October 2022. https://www.tghat.com/2023/01/12/government-of-tigray-bureau-of-agriculture-natural-resources-emergency-and-recovery-plan/

Thomas-Sharma, S., Abdurahman, A., Ali, S., Andrade-Piedra, J., Bao, S., Charkowski, A., Crook, D., Kadian, M., Kromann, P., et al. (2016). Seed degeneration in potato: the need for an integrated seed health strategy to mitigate the problem in developing countries. Plant Pathology, 65(1), 3–16. 10.1111/ppa.12439

Thomas-Sharma, S., Andrade-Piedra, J., Carvajal Yepes, M., Hernandez Nopsa, J., Jeger, M., Jones, R., Kromann, P., Legg, J. P., Yuen, J., & Forbes, G. (2017). A risk assessment framework for seed degeneration: Informing an integrated seed health strategy for vegetatively propagated crops. Phytopathology, 107(10), 1123–1135.

U.S. Environmental Protection Agency. 2009. USEPA: Expert Elicitation Task Force White Paper | US EPA Archive Document. https://archive.epa.gov/osa/pdfs/web/pdf/expert_elicitation_white_paper-january_06_2009.pdf (accessed 17 June 2025).

USAID. (2022). Strategic framework for early recovery, risk reduction, and resilience (ER4). https://www.usaid.gov/sites/default/files/2023-01/ER4_Framework-10.13.2022.pdf

USDA NIFA. (2018). Crop research technology readiness level (TRL).

van de Schoot, R., Depaoli, S., King, R., Kramer, B., Märtens, K., Tadesse, M. G., Vannucci, M., Gelman, A., Veen, D., Willemsen, J., and Yau, C. (2021). Bayesian statistics and modelling. Nat Rev Methods Primers, 1, 1. 10.1038/s43586-020-00001-2

Vasantha Raju, N., & Harinarayana, N. (2016). Online survey tools: A case study of Google Forms. National conference on scientific, computational & information research trends in engineering, GSSS-IETW, Mysore

White, R., Marzano, M., Fesenko, E., Inman, A., Jones, G., Agstner, B., and Mumford, R. (2022). Technology development for the early detection of plant pests: a framework for assessing Technology Readiness Levels (TRLs) in environmental science. Journal of Plant Diseases and Protection, 129, 1249–1261.

Wickham, H. (2016). ggplot2: Elegant Graphics for Data Analysis. Springer-Verlag New York, 3(2), 180–185. https://ggplot2.tidyverse.org

Wiebe, K., Zurek, M., Lord, S., Brzezina, N., Gabrielyan, G., Libertini, J., Loch, A., Thapa-Parajuli, R., Vervoort, J., & Westhoek, H. (2018). Scenario development and foresight analysis: exploring options to inform choices. Annual Review of Environment and Resources, 43, 545–570. 10.1146/annurev-environ-102017-030109

Williams, C. J., Wilson, K. J., and Wilson, N. (2021). A comparison of prior elicitation aggregation using the classical method and SHELF. J R Stat Soc Ser A Stat Soc, 184, 920–940. 10.1111/rssa.12691

Wittmann, M. E., Cooke, R. M., Rothlisberger, J. D., Rutherford, E. S., Zhang, H., Mason, D. M., and Lodge, D. M. (2015). Use of structured expert judgment to forecast invasions by bighead and silver carp in Lake Erie. Conservation Biology, 29, 187–197.

Wubaye, G. B., Gashaw, T., Worqlul, A. W., Dile, Y. T., Taye, M. T., Haileslassie, A., Zaitchik, B., Birhan, D. A., Adgo, E., & Mohammed, J. A. (2023). Trends in rainfall and temperature extremes in Ethiopia: Station and agro-ecological zone levels of analysis. Atmosphere, 14(3), 483. 10.3390/atmos14030483

Xing, Y., Hernandez Nopsa, J. F., Andersen, K. F., Andrade-Piedra, J. L., Beed, F. D., Blomme, G., Carvajal-Yepes, M., Coyne, D. L., Cuellar, W. J., Forbes, G. A., Kreuze, J. F., Kroschel, J., Kumar, P. L., Legg, J. P., Parker, M., Schulte-Geldermann, E., Sharma, K., & Garrett, K. A. (2020). Global cropland connectivity: A risk factor for invasion and saturation by emerging pathogens and pests. BioScience, 70(9), 744–758. 10.1093/biosci/biaa067

Yuen, J. E., and Hughes, G. (2002). Bayesian analysis of plant disease prediction. Plant Pathology, 1, 407–412. 10.1046/j.0032-0862.2002.00741.x

